# Evaluating effects of antibiotics across preclinical models of the human gastrointestinal microbiota

**DOI:** 10.1101/2025.01.23.634623

**Authors:** Thomas A. Auchtung, Armando I. Lerma, Keegan Schuchart, Jennifer M. Auchtung

## Abstract

While antibiotics play important roles in treating infections, disruption of the gastrointestinal microbiota during antibiotic treatment can lead to negative health consequences. However, for many antibiotics, the spectrum of activity has been determined for select isolates rather than for the range of microbes that populate the gastrointestinal tract. Here, we examined the response of communities of gastrointestinal microbes to antibiotics using two different model systems, human fecal minibioreactors and human microbiota associated mice. Communities established in minibioreactors using 12 different fecal donors were exposed to 12 different classes of antibiotics. Samples from three fecal donors were used to colonize germ-free mice from three different genetic backgrounds; progeny mice were then exposed to 6 of 12 antibiotics tested in minibioreactors. Initial bacterial community diversity was dependent on both the fecal donor and model system. Antibiotics affected a wide range of taxa across the phylogenetic spectrum, with many taxa similarly affected across treatments with different classes of antibiotics. Vancomycin, typically administered to treat Gram-positive bacterial infections, decreased the abundance of diverse taxa, including Gram-negative Bacteroidota species. Effects on some taxa were restricted by model system, indicating the importance of environmental context on antibiotic susceptibility. Altogether, these results indicate the complex interrelationships between microbiota composition and environmental context on antibiotic susceptibility and demonstrate strengths and weaknesses of each pre-clinical model system for evaluating effects of new antibiotics and other compounds with potential to disrupt the microbiota.

## INTRODUCTION

The microbes that colonize the gastrointestinal tract (GI microbiota) play key roles in many aspects of human physiology, including modulating activities of the digestive (Martinez-Guryn et al., 2018), endocrine (Chimerel et al., 2014), immune (Schluter et al., 2020), and nervous (Buffington et al., 2016) systems. While antibiotics play an essential role in the treatment of infections (Hutchings et al., 2019), antibiotics use can also alter the composition and function of the GI microbiota. Initial, high-throughput sequencing studies demonstrated the strong effect antibiotics can have on multiple taxa abundant in the gut microbiota (Dethlefsen et al., 2008; Dethlefsen and Relman, 2011). Since then, there have been studies on factors that affect antibiotic response (Raymond et al., 2016) or recovery (Ng et al., 2019; Palleja et al., 2018; Suez et al., 2018), or examined the functional changes in the microbiome (Zaura et al., 2015). Efforts are complicated by interindividual microbiome diversity, pre-existing conditions and exposures, treatment with more than a single drug, and fluctuations due to altered diet and lifestyle.

Traditional screening platforms for new antibiotics have focused on characterization of antimicrobial activity against a panel of select clinical isolates (Guay, 2007; Li et al., 2020), although new innovations continue to emerge that facilitate more robust screening approaches against larger panels of strains and microbial communities (Guzman-Rodriguez et al., 2018; Terekhov et al., 2018; Watterson et al., 2020). Studies conducted in mice (Ajami et al., 2018; Gu et al., 2020) and humans (Rashid et al., 2015; Raymond et al., 2016) typically have other objectives and rarely test multiple classes of antibiotics that would allow broader observations about the taxa affected across treatments. Tools that facilitate testing of new antibiotics against a larger panel of strains or microbial communities play an important role in rapidly identifying off-target effects on the commensal microbiota.

Recently, we reported effects of interindividual microbiota variation on susceptibility to colonization with the GI pathogen, *Clostridioides difficile* (Huang et al., 2025). As part of these studies, we characterized changes in microbiota composition in fecal communities from twelve healthy individuals in response to treatment with six different clinical antibiotics (Augmentin, azithromycin, cefaclor, ceftriaxone, clindamycin, and fidaxomicin) using fecal minibioreactor arrays (MBRAs), a continuous-flow culture model of the nutritional conditions of the lumen of the distal colon. In this work, we report an extension of these studies to investigate the effects of antibiotic treatment on microbiota composition using an additional panel of antibiotics (ciprofloxacin, doxycycline, imipenem, metronidazole, sulfamethoxazole, and vancomycin) tested across the same 12 fecal microbial communities, along with re-analysis of the initial published microbiota data (Huang et al., 2025) to facilitate comparisons between treatment with all twelve antibiotics. We also characterized the effects of treatment with six antibiotics (Augmentin, azithromycin, cefaclor, ceftriaxone, clindamycin, and fidaxomicin) on microbiota composition in human microbiota-associated mice (^HMA^mice) that were colonized with fecal microbial communities from 3 of 12 donors studied in the MBRA model. As expected, we observed that antibiotic treatment led to changes in microbiota composition across fecal samples and models. Overall, many Clostridiales species were highly susceptible to the antibiotics tested, whereas the classes of antibiotics that also inhibited Bacteroidales species were more limited. While some taxa responded similarly to antibiotics across the two experimental models, there were several notable differences, which could partially be due to differences in microbiota composition between models, but may also reflect differential responses of taxa to antibiotics between the models. In addition to taxa lost due to direct effects of antibiotics on susceptible microbes, we also observed a decrease in some taxa previously reported to be resistant to antibiotics, suggesting the importance of cross-feeding interactions that would cause additional impacts when disrupted.

## MATERIALS AND METHODS

### Fecal samples

As previously described (Huang et al., 2025), fecal samples were self-collected by 12 healthy individuals who had not been treated with oral antibiotics within the previous six months. Samples were collected from children aged 4-17 (n=3), adults aged 18-65 (n=12), and older adults aged >65 (n=2), with similar numbers of male and female participants. Studies were collected across the age span to capture differences that may be present, but were not specifically powered to evaluate differences in community composition by age. Adult participants provided consent to participate in the study. Children provided assent to participate along with parental consent to participate. Protocols for collection and use of fecal samples were reviewed and approved by Institutional Review Boards at Baylor College of Medicine (protocol number H-38014) and University of Nebraska-Lincoln (protocol number 18585). Details of fecal sample collection and cryopreservation can be found in our previously described study (Huang et al., 2025).

### Bioreactor experiments

Bioreactor experiments were performed in anoxic conditions at 37°C using minibioreactor arrays (MBRAs) as described previously (Auchtung et al., 2015). To prepare 1 L bioreactor medium (BRM3), a solution containing 1 g tryptone, 2 g proteose peptone number 3, 2 g yeast extract, 0.4 g sodium chloride, 1 g bovine bile, 5 mg hemin, 0.01 g magnesium sulfate, 0.01 g calcium chloride, and 2 ml Tween 80, was autoclaved at 121°C for 60 min. A mixture of 0.1 g arabinogalactan, 0.15 g maltose, 0.15 g D-cellobiose, 0.04 g D-glucose, 0.2 g inulin, 2 g sodium bicarbonate, 1 mg vitamin K1, 0.04 g potassium phosphate monobasic, and 0.04 g potassium phosphate dibasic was adjusted to pH 6.8, filter-sterilized, and added to the autoclaved solution.

Sources of reagents were previously described (Huang et al., 2025). Media and sterile MBRA were pre-reduced in an anaerobic chamber for >48 hours prior to use. Reactors were inoculated with fecal slurry to a final concentration of 1% w/v and allowed to equilibrate under anaerobic conditions for 16-18 hr. Media flow was initiated after equilibration at 1.875 mL/hr media (eight hour retention time). Communities were given 6 days to stabilize before treatment with antibiotics. Antibiotics were dosed every twelve hours for five days. Research grade antibiotics were obtained from the sources listed in Table A1 at the concentrations indicated in Table 1. Concentrations used for dosing were estimated from previously published measurements in human feces and/or bile (Brogard et al., 1982; Kager et al., 1981; Krook et al., 1981; Neu, 1974; Owen and Faragher, 1986; Pletz et al., 2004; Sears et al., 2012; Steinbakk et al., 1992; Sullivan, 2001), although fidaxomicin dosing was limited by its maximal solubility. Each fecal sample was tested with each antibiotic in duplicate, with the exception of cefaclor, ceftriaxone and untreated controls, which were tested four times for each fecal sample. Samples were collected for microbial community sequencing before antibiotic treatment (Day 0), after antibiotic treatment (Day 5), and two days following the end of antibiotic treatment (Day 7). Day 7 samples were limited to duplicate reactors from cefaclor, ceftriaxone and untreated communities. One mL samples were collected aseptically through the sampling port, cells were pelleted by centrifugation at ∼ 3000 X g for 5 min, and supernatants were transferred to new 96 well plates or 1.7 mL tubes prior to storage at -80°C. The heated anaerobic chamber (Coy laboratories) was maintained at 37°C with an atmosphere of 5% H2, 5% CO_2_, and 90% N_2_.

**Table 1.** Antibiotics tested in this study.

| Antibiotic | Class | MBRA<br>concentration<sup>1</sup> | <sup>HMA</sup>mice<br>concentration<sup>2</sup> |
| --- | --- | --- | --- |
| Augmentin | Penicillins | 25 µg/mL | 23.6 mg/kg |
| Azithromycin | Macrolides: Azalides | 400 µg/mL | 7.1 mg/kg |
| Cefaclor | Cephalosporins: 2<sup>nd</sup> generation | 67 µg/mL | 20.0 mg/kg |
| Ceftriaxone | Cephalosporins: 3<sup>rd</sup> generation | 200 µg/mL | 28.6 mg/kg |
| Ciprofloxacin | Fluoroquinolones | 175 µg/mL | ND |
| Clindamycin | Lincosamides | 150 µg/mL | 17.1 mg/kg |
| Doxycycline | Tetracyclines | 14.5 µg/mL | ND |
| Fidaxomicin | Macrolides: Tiacumins | 50 µg/mL | 5.7 mg/kg |
| Imipenem | Carbapenem | 9.5 µg/mL | ND |
| Metronidazole | Nitroimidazoles | 200 µg/mL | ND |
| Sulfamethoxazole | Sulfonamides | 12 µg/mL | ND |
| Vancomycin | Glycopeptides | 150 µg/mL | ND |

<sup>1</sup>MBRAs were treated twice daily for 5 days except as noted.
<sup>2HMA</sup>mice were treated once daily for 5 days.
ND, Not Determined

### Mouse experiments

Germ-free Swiss Webster mice were obtained from Baylor College of Medicine; germ-free C3H/HeN and C57BL/6J were obtained from the Nebraska Gnotobiotic Mouse facility. Fecal slurries from three human fecal samples (FS04, FS05, and FS06) were prepared under anaerobic conditions as previously described (Collins et al., 2015) and 100 μL were used to colonize each mouse intended for breeding (both breeding pairs and trios were used) under gnotobiotic conditions. Mice were then housed in individually ventilated cage racks under specific pathogen free conditions and were fed sterilized chow and water. All experiments were performed according to protocols approved by the Institutional Animal Care and Use Committees of Baylor College of Medicine (AN-6675) and University of Nebraska-Lincoln (IACUC #1668, #1680) and are in accordance with the NIH Office of Laboratory and Animal Welfare (OLAW) guidelines.

All experiments were performed with 6-10 week old mice, primarily from the F1 generation as indicated in Supplementary Table S1. Exceptions to testing in the F1 generation included Swiss Webster mice colonized with FS05, which were tested in the F3 generation, and Swiss Webster mice colonized with FS06 and C57Bl/6J mice colonized with FS04 and FS05, which were tested in the F2 generation. Swiss Webster mice colonized with FS05 were originally colonized at Baylor College of Medicine and imported to the University of Nebraska-Lincoln for these studies. Testing of other mice in the F2 generation was a result of limited fecundity of the original breeding pairs and trios (up to 5 per mouse background/fecal slurry). Previous studies had demonstrated that the microbiota composition stabilized following colonization of F1 generation with minimal compositional drift (Collins et al., 2015).

To analyze effects of antibiotics on microbiota composition, mice were administered 100 μl of research grade antibiotics (or sterile PBS) via orogastric gavage once daily for five days. To minimize the number of mice used in studies, both male and female mice were used, with animals housed 2-4/cage by sex. Concentrations of antibiotics used in mice were scaled based on doses reported for humans (factors used for scaling: 70 kg human, 25 g mouse, with doses adjusted by mouse body weight as indicated in Table 1). Table A2 provides the number of mice and their sex per treatment. Fecal samples were collected at the end of antibiotic treatment and used for microbial community sequencing analysis as described below.

### Growth of individual strains and testing of vancomycin susceptibility

Strains used in this study were originally isolated from one of the twelve fecal samples described above or were obtained from ATCC. For strains isolated from fecal samples, diluted fecal material was spread on non-selective agar (M2GSC (Miyazaki et al., 1997) or YCFA (Browne et al., 2016; Table A3) and incubated under anaerobic conditions for up to 72 hours. Colonies were passaged on culture media for one to two generations before cryopreservation of a stock in media supplemented with 7.5% DMSO or 15% glycerol. Partial taxonomic identity of strains was determined through sequencing of the V4 region of the 16S rRNA gene as described below. Strains isolated were tested for their ability to grow in medium with or without 150 μg/mL vancomycin. Colonies grown on YCFA were inoculated into BRM3 and YCFA in a 96-well polystyrene plate. Overnight cultures were inoculated 1:10 in fresh media, grown 2 hr, then diluted 1:30 into wells of fresh media +/-vancomycin. Growth at 37°C in the anaerobic chamber was monitored continuously by recording optical density at 600 nm in a Tecan Sunrise plate reader, with pre-read mixing on the low setting. Values are the average of two replicates grown in internal wells (to limit evaporation). Controls were uninoculated media and vancomycin-resistant strains *Enterococcus faecalis* and *Lactobacillus reuteri* (grown in MRS instead of YCFA).

*Akkermansia muciniphila* human-isolated strain BAA-835 and mouse-isolated strain YL44 were grown in YCFA, BRM3, or mixtures of components of BRM3 and YCFA (Table A4). Freezer stocks of the strains were struck on M2GSC and grown 2-3 days. Colonies were suspended in PBS and used to inoculate different medias. Four replicates of each culture condition were grown anaerobically in a 96-well polystyrene plate at 37°C. Aliquots of inoculum and 24 hr cultures were serially diluted and plated on M2GSC plates, which were incubated anaerobically at 37°C for 3 days before enumeration.

### Microbial community analysis through 16S rRNA gene sequencing

Bacteria were targeted for analysis because although protists, fungi, and archaea are also susceptible to some classes of antibiotics, typically only bacteria are abundant in the human GI tract. Frozen cell pellets (MBRA) or fecal samples (^HMA^mice) were initially disrupted by 0.1 mm bead-beating, followed by BioSprint 96 One-For-All Vet kit processing as described (Huang et al., 2025). DNA was amplified in duplicate with Phusion polymerase using Illumina barcoded primers 515F and 806R as described (Collins et al., 2015), then sequenced on a MiSeq using 2 x 250 kits according to manufacturers protocol by the investigators in the Nebraska Food for Health Center at the University of Nebraska-Lincoln. 16S rRNA gene data were deposited in NCBI’s Sequence Read Archive under BioProject ID PRJNA729569. Fastqs were processed by mothur 1.41.3, removing chimeras identified by uchime, mapping sequences against Silva release 132, and clustering OTUs at 99% identity using the OptiClust algorithm (Quast et al., 2013; Schloss et al., 2009). Because of the size of the study, two sequence processing runs were performed. The first run processed all sequences obtained from MBRAs. The second run processed all samples obtained from ^HMA^mice along with matched MBRA samples from the same fecal donors (FS04, FS05, and FS06) and replicate samples of the fecal inocula. For FS05, we sequenced one aliquot collected at the time of colonization of Swiss Webster mice and three replicates collected at the time of MBRA studies and colonization of ^HMA^ mice of other genetic backgrounds. For both studies, mothur version 1.48.1 was used to rarefy samples to 6944 reads prior to downstream analysis. OTU tables for each study are available in the Appendix (Mice, Supplementary Table S1; MBRA, Supplementary Table S2).

### Analysis of small organic acid levels by HPLC

We measured short and branched chain fatty acids from Day 5 samples collected from MBRA communities described above. A single sample was analyzed for each fecal donor/treatment combination, with the exception of untreated, cefaclor, and ceftriaxone, for which two replicates for each fecal donor/treatment were tested. Frozen supernatants were thawed and filtered through 0.22 μM filters, prior to injection of 100 μL into an Agilent 1260 Infinity LC system fitted with an HPX-87H column (BioRad) with a flow rate of 0.6 mL/min in 10 mM H_2_SO_4_ running buffer. The column was maintained at 55°C during the run. Peaks were detected by measuring absorption at 215 nm. The area under the curve was determined for every distinct peak present in ≥20% samples. Comparisons were made to standard curves of formate, acetate, propionate, butyrate, valerate, lactate, isovalerate, and isobutyrate resuspended in BRM3. Only acetate, butyrate, and isovalerate were detected in ≥20% of samples.

### Statistical testing

A combination of statistical packages were used for analysis including mothur version 1.48.1, R version 4.2.3, and GraphPad Prism version 10.2.3. R packages used included ANCOMBC vs 1.6.4, DESeq2 v 1.36.0, dplyr v 1.1.4, ggplot2 vs 3.5.1, microbiome 1.18.0, microViz 0.12.0, OTUtable 1.1.2, pairwise Adonis v 0.4.1, phyloseq v 1.40.0, rstatix v 0.7.2, stats v 4.2.3 tidyverse v 2.0.0, and vegan v 2.6-4. Microsoft Excel was also used for data management and calculation of averages as indicated. All code and files for analysis are at Zenodo: https://zenodo.org/records/14728946; doi:10.5281/zenodo.14728946.

For MBRA data, relative abundance of taxa at the OTU and phylum level was plotted using ATIMA, a Shiny-based interface for R packages for microbiome analysis available at atima.research.bcm.edu. The number of observed OTUs was determined using the estimate_richness command of phyloseq, means of replicate data were determined in Excel, data was plotted in GraphPad Prism and statistical significance of differences before and after treatment was determined by paired one-way ANOVA with Geisser-Greenhouse correction of sphericity of data and Holm-Sidak correction for multiple comparisons. Jaccard and Bray-Curtis dissimilarities were determined using mothur v 1.48.1, with a custom Excel template used to extract relevant data, which was plotted in GraphPad Prism. Statistical significance of differences from untreated communities was determined with paired one-way ANOVA with Geisser-Greenhouse correction of sphericity of data and Holm-Sidak correction for multiple comparisons. Differences in relative abundance of taxa in antibiotic-treated communities compared to untreated communities were identified using DESeq2 and ANCOM-BC2 at the genus and OTU levels. For genus-level analysis with DESeq2, OTU abundances were combined at the genus-level using the tax_glom function of phyloseq. Pairwise comparisons between each antibiotic-treated and untreated community was performed with DESeq2 with default parameters. For ANCOM-BC2, analysis was performed with the default parameters recommended in its tutorial (“ANCOM-BC2 Tutorial,” 2025) Log_2_ ratios of OTU abundance in antibiotic-treated and untreated communities were determined in Excel. Specifically, average abundances of each OTU across replicate samples for each fecal donor/treatment condition were determined. Log_2_ ratios between antibiotic-treated and untreated communities were then determined for each fecal sample and the mean of these log_2_ ratios across all fecal samples was determined. Ratios were matched with the list of genera (Supplementary Table S3) and OTUs (Supplementary Table S4) identified as significantly different following treatment with one or more antibiotics across analysis tools. Finally, the mean of log_2_ ratios across all fecal samples of genera determined to be differentially abundant were plotted as a heatmap using ggplot2. For tentative species-level assignments of 16S rRNA gene sequences, we used a combination of the Unassigner tool in BioConda and BLAST. As recently described (Tanes et al., 2024), the Unassigner tool aligns sequences to the Living Tree project reference database and determines the potential number of mismatches to the full length 16S rRNA gene sequence and the probability of accurate taxonomic assignment. We report taxonomy assigned with <4 mismatches and a p-value <0.1. In the case of more than one match, we report the top hit. In the case that high confidence taxonomic assignments could not be provided through Unassigner, BLAST (Altschul et al., 1990) against nr/nt database with default parameters excluding uncultured samples was used for tentative taxonomic assignment (100% minimum query coverage; 100% sequence identity). Representative sequences for each OTU are provided in Tables A6 to facilitate re-analysis as taxonomy continues to be refined, although we also acknowledge the limitations of partial 16S rRNA gene sequences for high confidence taxonomic assignments (Tanes et al., 2024). Similar procedures were used for analysis of ^HMA^mice data and comparisons between ^HMA^mice, fecal inocula, and bioreactor data. Compositional differences in untreated ^HMA^mice and fecal inocula were visualized through a combination of a heatmap of the 100 most abundant OTUs, which was generated using the plot_heatmap function of phyloseq, PCoA plots of Bray-Curtis dissimilarity calculated with vegan and plotted with ggplot2, and plotting of phylum level abundances, which were generated from OTU data using the tax_glom function in phyloseq, then plotted in GraphPad Prism. Statistical significance between treatment groups in the PCoA plot was determined with pairwise Adonis in R. Statistical significance of differences in phyla abundance was determined with one-way ANOVA with Brown-Forsythe and Welch correction for unequal variance and Dunnett’s correction for multiple testing in GraphPad Prism. The number of observed OTUs and Bray-Curtis and Jaccard dissimilarities were determined using mothur v.1.48.1 with Microsoft and plotted in GraphPad Prism, with statistical tests performed as described above. Species with differential abundance between treated and untreated communities on Day 5 were determined through a combination of ANCOM-BC2 and DESeq2 at the genus and OTU levels as described above. The average log2 fold changes from untreated communities were calculated (Supplementary Table S5, differentially abundant genera; Supplementary Table S6, differentially abundant OTUs) and the data was visualized through a heatmap as described above. Comparisons between MBRA and ^HMA^mice models were performed at the OTU level (observed OTUs and Bray-Curtis dissimilarity) and the genus-level (differentially abundant taxa; Supplementary Table S7). For differentially abundant taxa shared between models, taxa were matched at the genus level and the relative abundance of taxa in antibiotic-treated samples compared to untreated controls was plotted for both models when at least one model had statistically significant values; the heatmap was plotted in GraphPad Prism.

## RESULTS

### Magnitude of microbiota disruption varies by fecal donor and antibiotic class in fecal MBRAs

Independent minibioreactor array (MBRA) communities were established from the stool of twelve healthy human donors (FS01 - FS12). After allowing one week for communities to stabilize (Auchtung et al., 2015), analysis of microbial communities through sequencing the V4 region of the 16S rRNA gene revealed that communities sampled before antibiotic treatment (Day 0) varied in diversity, with a median of between 108 and 178 observed operational taxonomic units (OTUs; clustered at 99% nucleotide identity) per donor and a mean relative abundance of 45.0% Bacteroidota, 33.5% Bacillota, 18.3% Pseudomonadota, 1.8% Verrucomicrobiota, 0.8% Fusobacteriota, 0.3% Synergistota, 0.1% Cyanobacteriota, and 0.08% Actinomycetota (Figure A1). These communities represent a subset of the starting inoculum (∼50% reduction in taxonomic diversity)(Huang et al., 2025), similar to what has been reported for other *in vitro* and animal models of the GI microbiome (Aluthge et al., 2020; Collins et al., 2015; Rajilić-Stojanović et al., 2010; Van Den Abbeele et al., 2013, 2010). We examined the response of these communities to five days of twice-daily treatment with twelve different clinically relevant antibiotics (Table 1) or an equivalent dose of water for untreated controls. Each donor/treatment combination was tested twice, with the exception of cefaclor, ceftriaxone and untreated samples, which were tested four times. Changes in microbial community composition were assessed by 16S rRNA gene sequencing at the end of antibiotics (Day 5).

Overall, we observed a range of responses to different antibiotics across all fecal samples (Figure 1). Treatment with all antibiotics except sulfamethoxazole led to significant declines in observed OTUs between samples collected before and after antibiotics (Figure 1A). Levels of observed OTUs were not significantly different following treatment with water in untreated controls. Observations for Jaccard dissimilarity, which assesses compositional similarity based upon presence of shared OTUs, were similar to those for observed OTUs, with all antibiotics with the exception of sulfamethoxazole and azithromycin showing significantly larger changes in Jaccard dissimilarity compared to untreated communities (Figure 1B). Fewer antibiotics showed significant changes in Bray-Curtis dissimilarity, which assesses changes in community structure based upon abundances of shared OTUs (Figure 1C). The extent to which antibiotic treatment led to loss of OTU richness also varied across fecal donors (Figure 1D), with all fecal communities except for fecal donor #6 (FS06) exhibiting statistically significant decreases in OTU richness. However, decreases in median richness in response to all antibiotics varied from 15% (7-27% interquartile range (IQR)) for FS06 to 32% (21-43% IQR) for FS02, pointing to donor-specific responses to antibiotic treatment.

**Figure 1.**
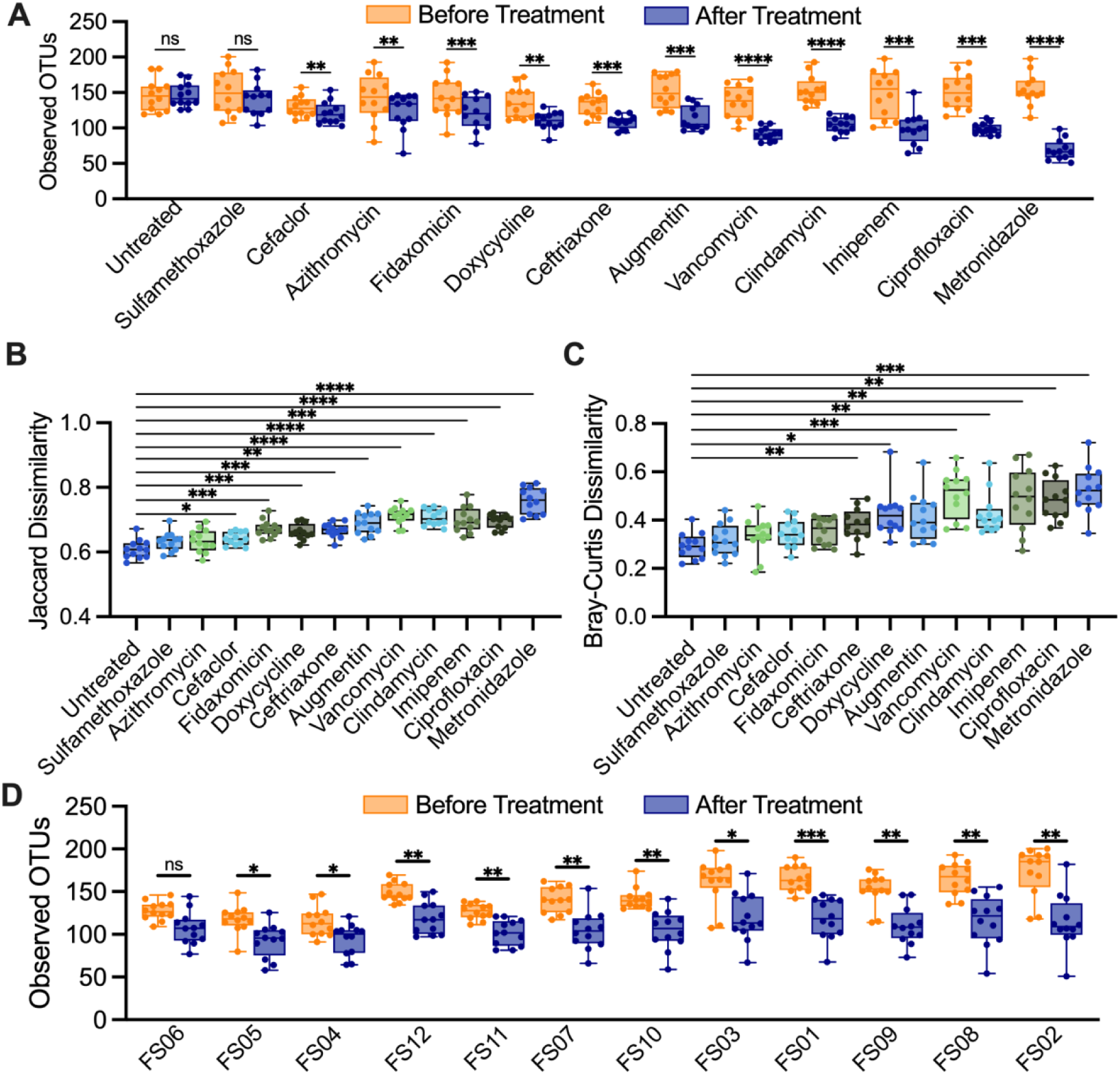
Effects of antibiotic treatment on microbial diversity for human fecal communities cultured in MBRAs. (A) Observed OTUs before and after treatment with the treatment indicated on the x-axis. (B) Jaccard and (C) Bray-Curtis dissimilarity of samples before and after treatment with the treatment indicated on the x-axis. (D) Observed OTUs before and after treatment with antibiotics for each of the twelve fecal sample MBRA communities as indicated on the x-axis. For all panels, individual points were plotted for the mean of replicate samples for each of the 12 fecal communities tested. Further, median values were plotted as horizontal lines, interquartile (IQR) ranges were plotted as boxes, and range of data was plotted as whiskers. Statistical significance was determined by paired ANOVA with Geisser-Greenhouse correction for unequal variability of differences and Holm-Šidák correction for multiple comparisons. In (A) and (D), comparisons were performed between samples before and after treatment. In (B) and (C), all treatments were compared to untreated communities. ns>0.05. *, p<0.05; **, p<0.01; ***, p<0.001, ****; comparisons with p>0.05 in panels (B) and (C) were not shown.

We also examined the persistence of microbial disruption after antibiotic treatment by assessing community composition two days following the end of antibiotic treatment in two replicates per treatment/donor (Figure A2; Recovery). While OTU richness remained significantly lower than before treatment for most antibiotics tested, a subset of antibiotics showed partial recovery of OTU richness (Figure A2A; imipenem, ciprofloxacin, metronidazole), microbiota composition (Figure A2B; Augmentin, imipenem, ciprofloxacin, metronidazole) and/or microbiota structure (Figure A2C; vancomycin, imipenem, ciprofloxacin, metronidazole) two days following cessation of antibiotics.

### Conserved effects of antibiotic treatments on taxonomic diversity in fecal MBRAs

We next assessed how antibiotic treatment affected levels of specific taxa across fecal communities cultured in MBRAs using a combination of ANCOM-BC2 (Lin and Peddada, 2024) and DESeq2 (Love et al., 2014). We performed analyses primarily at the genus level, as the number of genera shared in ≥ 50% of fecal sample communities after treatment (n=26 genera) was higher than the number of OTUs shared in ≥ 50% of fecal sample communities after treatment (n=23 OTUs). We observed a total of 49 genera were differentially abundant following treatment with one or more antibiotics in one or more tests (Figure 2). As had been reported previously (Nearing et al., 2022), ANCOM-BC2 identified a smaller number of genera with significant differences from untreated communities across antibiotics (n=14) compared to DESeq2 (n=46), although 11 genera were identified as having significant differences from untreated communities following treatment with one or more antibiotics in both tests (Figure 2, black circles; Supplementary Table S3).

**Figure 2.**
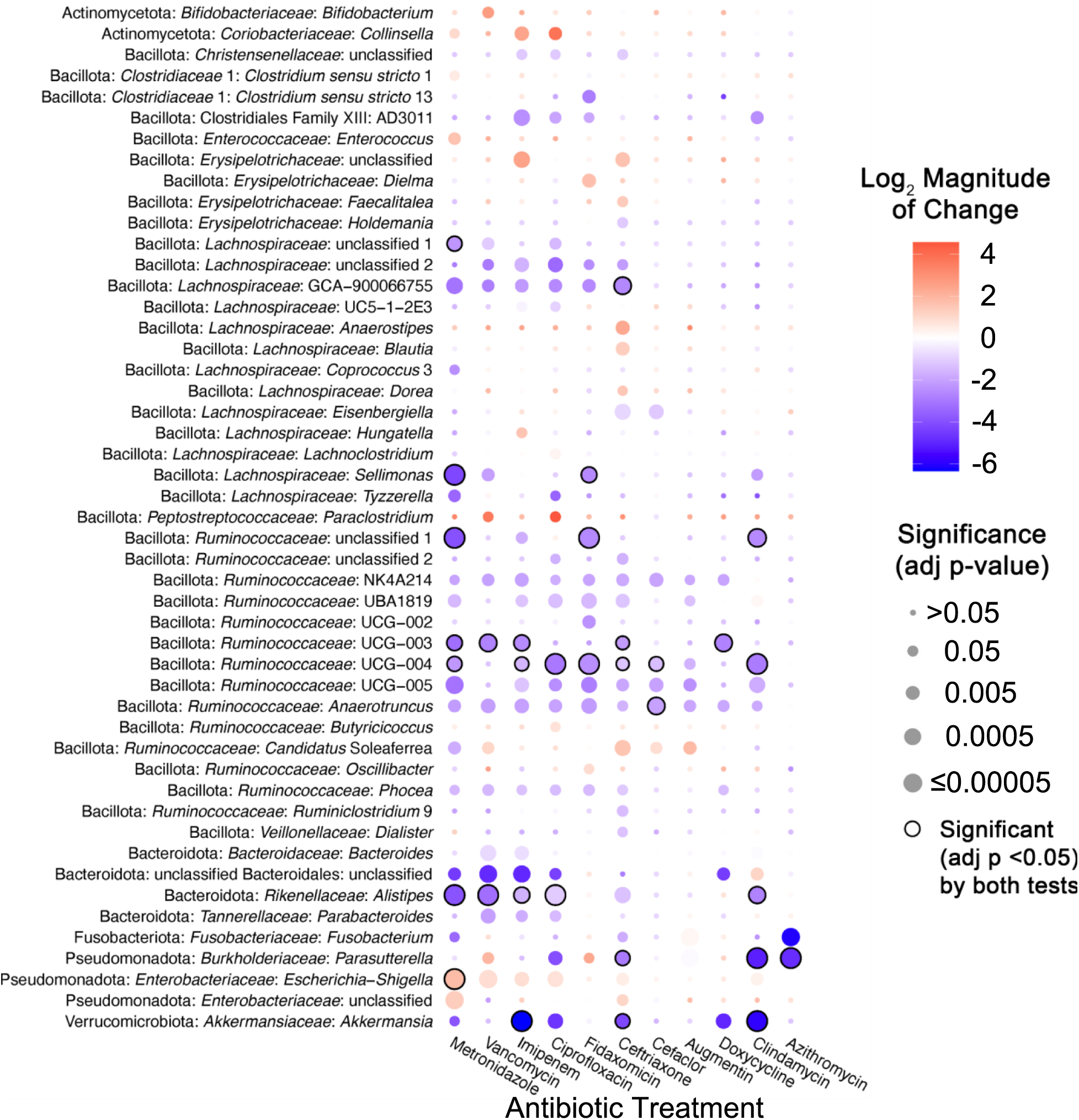
Taxa in bioreactor communities affected by different antibiotics. Heat map of the statistically significant changes in abundance of genera between antibiotic-treated and untreated communities. Shading represents the magnitude of the log2 mean ratio between all untreated communities and communities treated with the indicated antibiotic. The size of the circle corresponds to statistical significance, which was determined using with DESeq2 unless otherwise noted in Supplementary Table S3. Black lines around circles indicate taxa identified as significant by both DESeq2 and ANCOM-BC2. Only genera with a significant change for at least one antibiotic (FDR-corrected p<0.05) were plotted. There were no genera whose abundance changed significantly in Sulfamethoxazole-treated communities compared to untreated communities.

There was a wide range in the effects of different antibiotics on these 49 differentially abundant genera, with each antibiotic demonstrating a unique profile of affected taxa (Figure 2). The largest number of differentially abundant genera were observed in communities treated with ceftriaxone (27 genera), whereas there were no genera whose abundance differed significantly from untreated controls in sulfamethoxazole-treated communities. Overall, there was a strong correlation (R^2^=0.52, p=0.0081) between lower levels of observed OTUs following antibiotic treatment and the number of genera that were differentially abundant in communities treated with the same antibiotic (Figure A3). Notable exceptions included fidaxomicin and metronidazole-treated communities, which had higher and lower levels of genera that were significantly different from untreated communities than would be predicted by the changes in observed OTUs.

The majority of affected genera were members of Bacillota phylum (38 genera) including the *Ruminococcaceae* (14 genera), and *Lachnospiraceae* (13 genera) families, which are consistent with the prevalence of these families in fecal samples and bioreactor microbial communities, as well as the diversity of genera within these families (Figure A1; Supplementary Table S2). Four genera within the Bacteroidota (*Alistipes, Bacteroides, Parabacteroides,* unclassified Bacteroidales genus) showed significant differences in one or more antibiotic-treated communities compared to untreated communities. Identification of a single *Bacteroides* genus in the *Bacteroidaceae* family as differentially abundant reflects the more limited diversity of *Bacteroidaceae* genera in these communities, as similar numbers of OTUs classified as *Bacteroidaceae* (18 OTUs) and *Ruminococcaceae* (17 OTUs) were differentially abundant compared to untreated communities (Supplementary Table S4). Notable changes for members of other phyla include significantly lower levels of *Akkermansia* (Verrucomicrobiota) and higher levels of *Escherichia-Shigella* (Pseudomonadota) in communities treated with six different antibiotics. There were also eight abundant genera (total relative abundance across all samples >0.5%) that were not significantly different between untreated communities and communities treated with antibiotics (Supplementary Table S8). These include *Erysipelatoclostridium, Phascolarctobacterium, Bilophila, Lachnospiraceae* genus UCG-004, *Paeniclostridium, Erysipelotrichaceae* genus UCD-003, *Flavonifractor,* and *Agathobacter*.

In total, 27 genera were significantly lower in one or more antibiotic-treated communities, 17 genera were significantly higher in communities treated with one or more antibiotics, and 5 genera were higher or lower than untreated communities depending upon the antibiotic used for treatment. Some taxa were affected similarly across many classes of antibiotics. Fifteen genera were significantly different in communities treated with five or more antibiotics compared to untreated, including ten members of the Bacillota phylum: *Ruminococcaceae* genera *Anaerotruncus, Candidatus* Soleaferrea, NK4A214, UCG-003, UCG-004, UCG-005, UBA1819, and *Phocea*, and *Lachnospiraceae* GCA-900066755 and an unclassified genus. An unclassified Bacteroidales genus (Bacteroidota), *Alistipes* (Bacteroidota), *Akkermansia* (Verrucomicrobiota), and two Pseudomonadota, *Parasutterella* and *Escherichia-Shigella* were also significantly different in communities treated with five or more antibiotics compared to untreated.

### Vancomycin inhibits growth of Bacteroidota isolates in pure culture

Treatment with vancomycin led to a decrease in 9 genera of Bacillota as well as 4 genera of Bacteroidota. At the OTU level, 13 OTUs classified as Bacillota and 18 OTUs classified as Bacteroidota (including 14 *Bacteroides* species) were significantly lower in vancomycin-treated communities compared to untreated communities (Supplementary Table S4). The limited number of studies documenting the sensitivity of gastrointestinal Bacteroidota isolates to vancomycin (Citron et al., 2012; Ednie et al., 2003; Yehya et al., 2013) were done under different conditions and concentrations of vancomycin. The 150 μg/mL used in these studies are higher than typical fecal concentrations when vancomycin is administered intravenously (Currie and Lemos-Filho, 2004) but lower than when taken orally (Gonzales et al., 2010; Thabit and Nicolau, 2015). To further examine the sensitivity of Bacteroidota species to vancomycin, we tested sensitivity of multiple type strains and fecal isolates to 150 μg/mL vancomycin when grown in bioreactor media (BRM3) and a broad range growth media for gut microbial isolates (YCFA, (Browne et al., 2016)). This study included eight *Bacteroides*, two *Phocaeicola*, two *Parabacteroides*, and one *Alistipes* (Figure A4). One strain, *Bacteroides cellulolyticus*, grew poorly in BRM3. None of the Bacteroidota strains were able to grow in the presence of 150 μg/mL vancomycin in BRM3 or YCFA media. Control strains of Gram-positive *Enterococcus faecalis* and *Limosilactobacillus reuteri* and Gram-negative *Escherichia coli* were also unable to grow at this level of vancomycin when in BRM3, but were resistant in YCFA (or MRS for *L. reuteri*).

### Magnitude of microbiota disruption varies by fecal donor and mouse background in ^HMA^mice

To determine whether the effects of antibiotic treatment on microbial diversity were similar in another commonly used model of the human GI microbiome, we colonized germ free mice from three genetic backgrounds (C57Bl/6J, C3H/HeN, and Swiss Webster) with fecal samples 4, 5, and 6. Previously, we had used C57Bl/6J mice stably colonized with fecal material pooled from twelve healthy humans to investigate susceptibility to *C. difficile* infection (Collins, 2015; Auchtung, 2020). In these previous studies, mice were initially colonized with fecal material under germ-free conditions, then bred under specific pathogen free conditions. The composition of the fecal microbiota in originally colonized germ-free mice (P1 generation) was most similar to the fecal inoculum, with a shift in microbiota composition observed in the F1 generation that was stable in subsequent generations (F2 and greater). We used the same approach here to colonize mice with fecal material under germ-free conditions and test the effects of antibiotic treatment in progeny mice. Progeny mice were treated once daily by orogastric gavage for five days with one of six different antibiotics (Augmentin, azithromycin, fidaxomicin, cefaclor, ceftriaxone, or clindamycin) or PBS for untreated controls (n=4-12 mice/treatment; Table A2). Antibiotic concentrations are listed in Table 1. Fecal samples were collected from mice at the end of treatment and microbiota community composition was analyzed through sequencing of the V4 region of the 16S rRNA gene. Analysis of untreated mice demonstrated that both mouse strain and fecal donor influenced taxonomic composition and overall diversity. Differences in taxonomic composition were primarily driven by the fecal donor used for colonization (Figure 3A, Figure A5). However, there were significant differences in microbiota richness (Figure 3B) and microbial community structure (Figure A6) within mice of different genetic backgrounds colonized by samples from a single fecal donor. Overall, mouse communities had a mean relative abundance of 58.0% Bacteroidota, 36.0% Bacillota, 2.9% Verrucomicrobiota, 1.9% Pseudomonadota, 0.8% Fusobacteriota, 0.2% Actinomycetota, 0.2% Mycoplasmatota, and 0.03% Cyanobacteriota (Figure 3C-3E). Mice colonized with FS05 had significantly lower numbers of total OTUs, with a mean of 141.5 OTUs/sample, compared to mice colonized with FS04 (mean of 165.3 OTUs) or FS06 (mean of 165.1 OTUs; Figure A7A). C3H/HeN mice also exhibited significantly lower levels of observed OTUs, with a mean of 133.4 OTUs/sample, compared to 167.5 for C57Bl/6J and 171.7 for Swiss Webster (Figure A7B).

**Figure 3.**
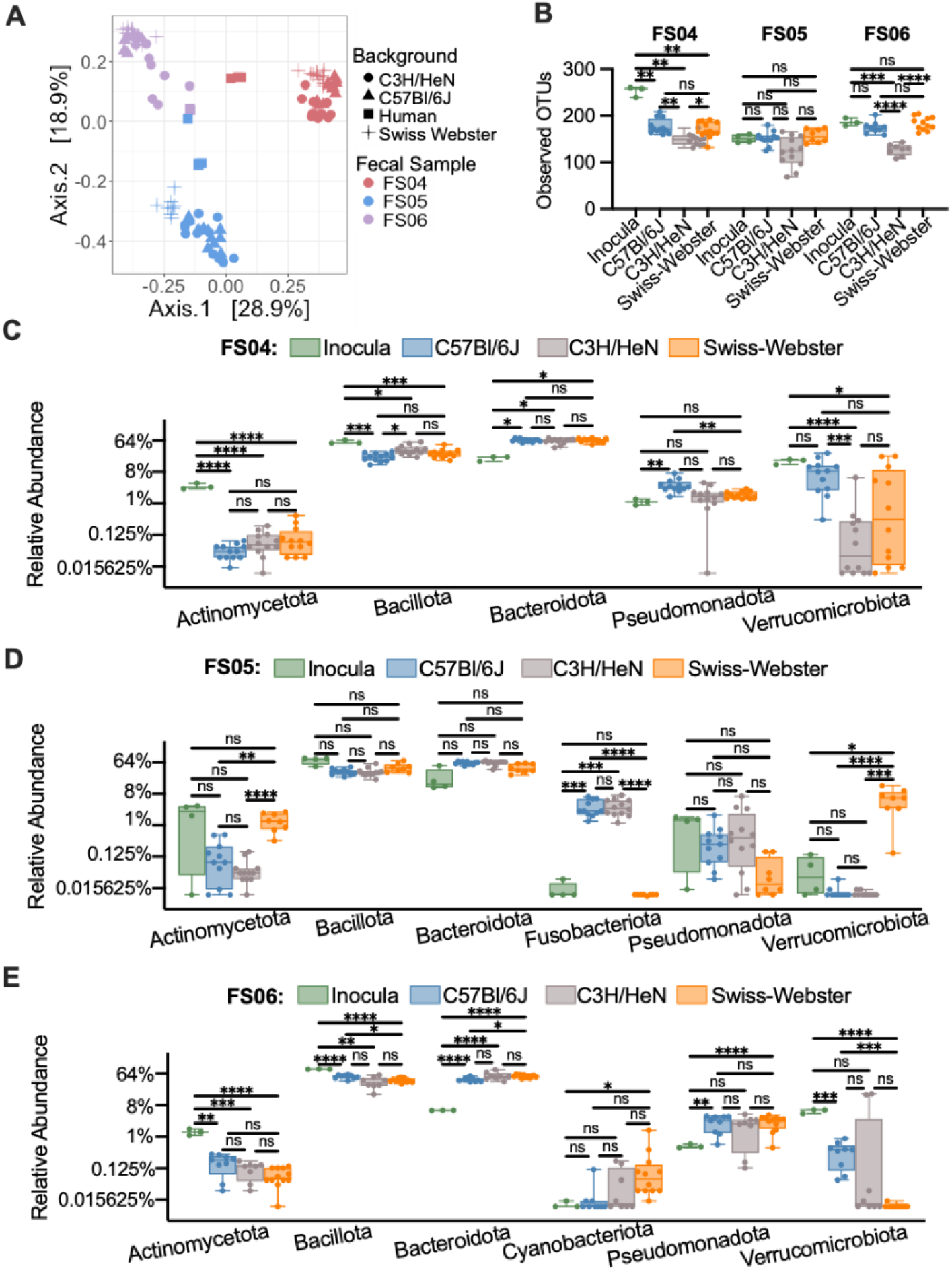
Comparison of microbiota community composition and structure between human fecal inocula and untreated ^HMA^mice. (A) PCoA plot of Bray-Curtis dissimilarities between replicate samples of fecal inocula and untreated ^HMA^mice. Each individual point represents a unique sample. Fecal donor (stool) is indicated by color and background (fecal inocula or mouse genetic background) is indicated by shape. Pairwise ANOVA indicated statistically significant separation by fecal donor and background, with the exception of comparisons between Swiss Webster and C57Bl/6J mice (Table A5). (B) Observed OTUs plotted by fecal donor and model. Each point represents an individual sample, with cross bars indicating medians, boxes indicating IQR, and whiskers indicate range of data. In (C)-(E), relative abundance of phyla across fecal inocula and untreated ^HMA^mice colonized with (C) FS04, (D) FS05, and (E) FS06. In (B)-(E), statistical significance of differences within a fecal sample were determined by one-way ANOVA with Brown-Forsythe correction for unequal variances and Dunnett’s T3 correction for multiple comparisons. *, p<0.05; **, p<0.01; ***, p<0.001; ****, p<0.0001; ns, p>0.05.

To determine the effects of antibiotic treatment on bacterial richness, we compared differences between antibiotic-treated and control (PBS-treated) mice. Across all mice tested, we observed that treatment with all antibiotics with the exception of cefaclor led to significantly lower levels of observed OTUs compared to untreated (Figure A8). The extent to which treatment with other antibiotics impacted bacterial richness varied by fecal community and mouse background (Figure 4). Ceftriaxone, fidaxomicin, azithromycin, and clindamycin treatment all led to significantly lower levels of OTUs overall in mice colonized with FS04 (Figure 4A) and FS06 (Figure 4E), whereas levels of observed OTUs were not significantly lower than in untreated mice following treatment with ceftriaxone and fidaxomicin in mice colonized with FS05 (Figure 4C). Comparing across mice of different genetic backgrounds colonized with the same fecal sample (Figures 4B, 4D, and 4F), it is clear that both mouse genetic background and fecal community effect the response to antibiotic treatment. Clindamycin resulted in the most significant differences in richness compared to untreated mice across all mice and fecal communities tested.

**Figure 4.**
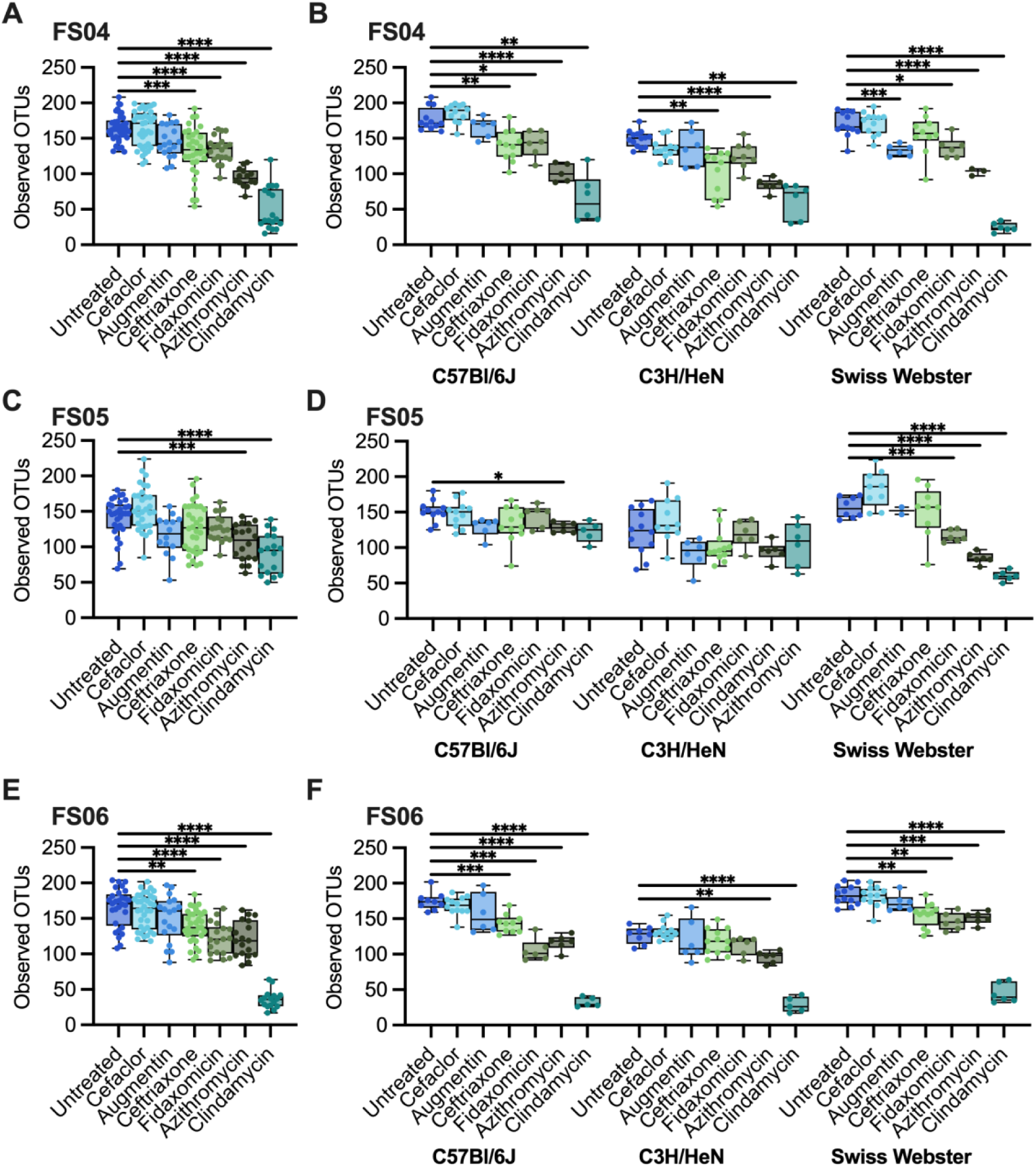
Changes in abundance of Observed OTUs following treatment with antibiotics. Levels of observed OTUs measured in replicate ^HMA^mice after five days of treatment with the indicated antibiotic or mock-treatment with PBS. Samples are presented for mice colonized with (A,B) FS04, (C,D) FS05, and (E,F) FS06. In panels (A), (C), and (E) data are plotted for all mice colonized with the same fecal sample without respect to mouse genetic background. In panels (B), (D), and (F), data are separated by genetic background. Each point represents an individual sample, with cross bars indicating medians, boxes indicating IQR, and whiskers indicate range of data. Statistical significance of differences relative to untreated mice were determined by one-way ANOVA with Brown-Forsythe correction for unequal variances and Dunnett’s T3 correction for multiple comparisons. *, p<0.05; **, p<0.01; ***, p<0.001; ****, p<0.0001; ns, p>0.05.

### Conserved effects of antibiotic treatments on taxonomic diversity in ^HMA^mice

We examined how antibiotics altered taxonomic representation in ^HMA^mice using the approach described above for MBRA communities. Overall, a total of 78 genera were identified as differing significantly between one or more antibiotic-treated groups and the untreated group. Again, DSeq2 testing identified a larger number of genera as significantly different between antibiotic-treated and untreated communities (77 genera) compared to ANCOM-BC2 (51 genera), although there was higher consensus between the two tests (49 genera; Figure 5, circled comparisons; Supplementary Table S5).

**Figure 5.**
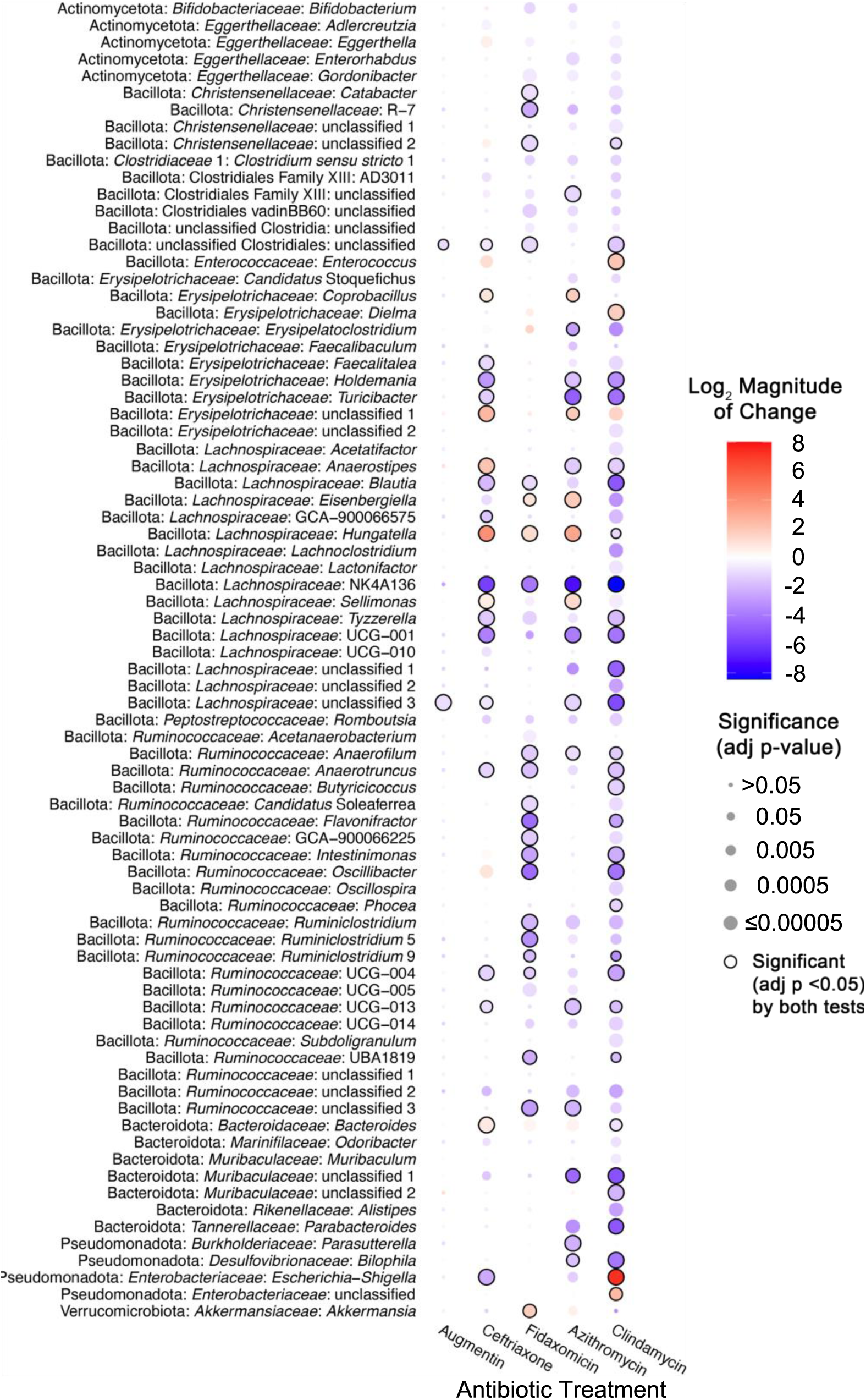
Taxa in ^HMA^mice affected by different antibiotics. Heat map of the statistically significant changes in abundance of genera between antibiotic-treated and untreated ^HMA^mice. Shading represents the magnitude of the log2 mean ratio between all untreated communities and communities treated with the indicated antibiotic. The size of the circle corresponds to statistical significance, which was determined using with DESeq2 unless otherwise noted in Supplementary Table S5. A black line around the circle indicates significance by both DESeq2 and ANCOM-BC2. Only genera with a significant change for at least one antibiotic (FDR-corrected p<0.05) were plotted. There were no genera whose abundance changed significantly in cefaclor-treated communities compared to untreated communities.

Each antibiotic demonstrated a unique profile of affected taxa (Figure 5), with the largest number of differentially abundant genera observed in mice treated with clindamycin (71 genera), whereas there were no genera whose abundance differed significantly from untreated mice in cefaclor-treated mice. There was a stronger correlation (R^2^=0.835, p=0.0011) between lower levels of observed OTUs following antibiotic treatment and a higher number of genera that were differentially abundant in communities treated with the same antibiotic (Figure A9), with Augmentin-treated mice the only notable exception, with lower levels of genera that were significantly different from untreated communities than would be predicted by the level of observed OTUs following treatment.

Genera that were differentially abundant in antibiotic-treated mouse communities compared to untreated controls belong to five different phyla (Actinomycetota, Bacillota, Bacteroidota, Pseudomonadota, and Verrucomicrobiota). Again, the majority of affected genera were members of the Bacillota phylum (61 genera) including the *Ruminococcaceae* (23 genera), *Lachnospiraceae* (16 genera), and *Erysipelotrichaceae* (10 genera) families, consistent with the prevalence of these families in ^HMA^mice (Figure 3; Supplementary Table S1). Seven genera within the *Bacteroidota* showed significant differences compared to untreated communities*: Bacteroides, Alistipes, Parabacteroides*, *Odoribacter*, and three genera from the *Muribaculaceae* family; as with MBRA communities, higher numbers of *Bacteroides* OTUs were observed as differentially abundant compared to genera (Supplementary Table S6). Notable changes for members of other phyla were significantly higher levels of *Akkermansia*(Verrucomicrobiota) in communities treated with fidaxomicin and azithromycin, loss of *Bifidobacterium, Adlercreutzia, Enterorhabdus,* and *Gordonibacter* (all Actinomycetota) in two or more antibiotic treated communities, and alterations in the levels of four *Pseudomonadota* genera: *Parasutterella*, *Bilophila*, *Escherichia-Shigella*, and an unclassified *Enterobacteriaceae* genus. There were also two abundant genera (*Phascolarctobacterium* and *Fusobacteria*) that were not significantly different between untreated communities and communities treated with antibiotics.

In total, 60 genera were only significantly lower in one or more antibiotic-treated communities. Ten of these genera were significantly lower in mice treated with four different classes of antibiotics compared to untreated mice, including those in *Lachnospiraceae* (*Blautia*, NK4A136, UCG-001, an unclassified genus, and *Tyzzerella*), *Ruminococcaceae* (*Anaerotruncus* and UCG-004), *Romboutsia* (*Peptostreptococcaceae*), and Clostridiales (a family XIII genus and an unclassified genus). In total, six genera were only significantly higher in one or more antibiotic-treated communities. For example, an unclassified *Erysipelotrichaceae* was increased for three different antibiotic treatments. In total, twelve genera were significantly higher or lower than untreated communities, depending upon the antibiotic used for treatment. For example, *Escherichia-Shigella* was significantly higher in clindamycin-treated mice, but significantly lower in azithromycin and ceftriaxone-treated mice.

### Comparison of effects of antibiotic treatment between MBRA and ^HMA^mice

When we compared between the subset of antibiotics tested in both models, we observed that most of the antibiotics tested led to reduction in levels of observed OTUs in antibiotic-treated communities compared to untreated controls across both models (Figure 6A). The two antibiotics that did not follow this trend were cefaclor, which led to significantly lower levels of observed OTUs in MBRAs but not in ^HMA^mice, and azithromycin, which led to significantly lower levels of observed OTUs in ^HMA^mice but not in MBRAs. There were no significant differences in microbiota structure between untreated and antibiotic-treated communities in the MBRA model, whereas clindamycin and azithromycin treatment led to significant changes in community composition compared to untreated controls (Figure 6B). In total, 88 genera were identified as significantly different in MBRA (Figure 2, 49 genera) and/or ^HMA^mice (Figure 5, 79 genera) treated with one or more antibiotics compared to untreated controls. Of these 88 genera, we focused on comparisons of the 57 genera that were present in ten or more samples of ^HMA^mice and MBRA communities to facilitate robust comparisons. Comparing these shared genera demonstrated 22 statistically significant differences in antibiotic-treated communities compared to controls that were similar across both models (Figure 6C, black boxes), as well as 5 statistically significant differences in antibiotic-treated communities that differed between models (Figure 6C, yellow boxes). Significant differences that were shared between models included lower levels of multiple *Ruminococcaceae* and *Lachnospiraceae* genera in response to treatment with ceftriaxone, clindamycin, and/or fidaxomicin, decreases in *Alistipes* in response to clindamycin-treatment, decreases in *Parasutterella* in azithromycin-treated communities, increases in the levels of unclassified *Erysipelotrichaceae* and *Enterobacteriaceae* genera in ceftriaxone-treated communities, increases in levels of *Escherichia-Shigella* in clindamycin-treated communities, and increases in *Dielma* in fidaxomicin-treated communities. For those genera/antibiotic combinations that were statistically significant in one model (Figure 6C starred) and not the other (Figure 6C unstarred)(n=166), 68% demonstrated a similar, non-significant change in taxa abundance in the other model.

**Figure 6.**
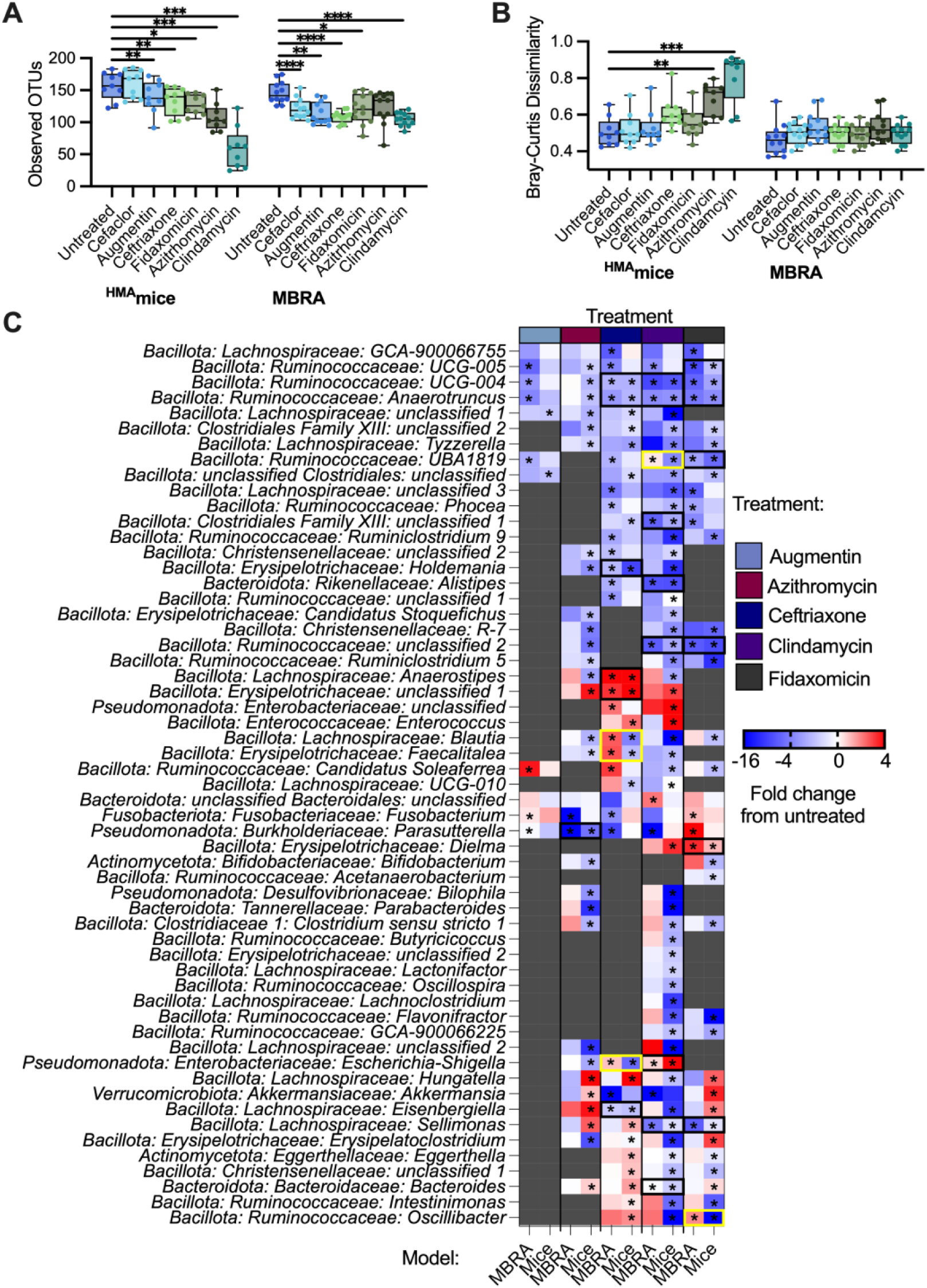
Similarities and differences in responses to antibiotics between ^HMA^mice and MBRA communities. (A) Observed OTUs after treatment with the antibiotics indicated on the x-axis. Individual points were plotted from the mean of replicates for one of the nine mouse genetic background/fecal donor combinations tested of from one of the twelve fecal samples tested in MBRA. Statistical significance of difference from untreated within each model was determined by paired ANOVA with Geisser-Greenhouse correction for unequal variability of differences and Holm-Šidák correction for multiple comparisons. (B) Mean Bray-Curtis dissimilarities of antibiotic-treated communities compared to untreated communities. Within group variation among untreated communities is plotted as a control for ^HMA^mice and MBRA communities. As in (A), individual points represents means calculated across replicates for nine mouse genetic background/fecal donor combinations or twelve MBRA fecal microbial communities. Statistical significance was determined by one-way ANOVA with Brown-Forsythe correction for unequal variances and Dunnett’s T3 correction for multiple comparisons. In (A) and (B), median values were plotted as horizontal lines, interquartile (IQR) ranges were plotted as boxes, and range of data was plotted as whiskers. *, p<0.05; **, p<0.01; ***, p<0.001, ****; comparisons with p>0.05 were not shown. (C) Heat map of the genera with statistically significant changes in antibiotic-treated and untreated communities shared between ^HMA^mice and MBRA. Shading represents the magnitude of fold change between all untreated communities and communities treated with the indicated antibiotic. Model tested is indicated in the x-axis, with antibiotics tested indicated across the top. Genus level classifications are provided on the y-axis. Abundance data was plotted for both models for a genus if that genus was statistically significant in at least one of the models in the antibiotic tested. Statistical significance of the fold change is indicated by an asterisk. Data are replotted from Figure 2 and 5 to facilitate direct comparisons.

## DISCUSSION

Here we examined the effect of multiple antibiotics on the communities of human fecal bacteria established in two different model systems. As expected, we observed changes in bacterial richness and community structure, along with changes in the abundance of specific taxa compared to untreated controls for most antibiotics tested. Overall, we observed that abundances of many genera of Gram-positive anaerobes in the Bacillota phylum (primarily members of the *Lachnospiraceae* and *Ruminococcaceae* families) were lower in communities treated with most of the antibiotics tested, although there were some variations in responses between models. We also observed differences in the abundance of multiple genera in the Gram-negative phylum Bacteroidota following treatment with several classes of antibiotics. Consistent with previous studies (Looft and Allen, 2012), we observed higher levels of potential pathobiont *Escherichia-Shigella* in communities treated with several classes of antibiotics. *Enterococcus*, another potential pathobiont demonstrated to increase following antibiotic treatment (Dubin and Pamer, 2017), was also significantly higher in metronidazole-treated MBRA communities and ceftriaxone and clindamycin-treated mice. *Akkermansia* were present at significantly lower levels in MBRA communities treated with ceftriaxone, ciprofloxacin, fidaxomicin, imipenem, and metronidazole. Finally, multiple Actinomycetota genera in the class *Eggerthellaceae* were lower in ^HMA^mice treated with azithromycin, clindamycin, and fidaxomicin.

While we observed several taxa that were affected similarly by antibiotic treatment across models, we observed many differences. There are many factors known to play a role in susceptibility to antibiotics, including the composition of phage populations (Modi et al., 2013), bacterial growth rate (Greulich et al., 2015), the presence of mucin (Samad et al., 2019), biofilms (Sharma et al., 2019), immune cells (Ramiro et al., 2016), and pharmacodynamics (Onufrak et al., 2016). However, another major difference between the models was the abundance of taxa prior to treatment. A direct comparison of untreated ^HMA^mice, MBRA communities before antibiotic treatment, and the fecal inocula used to colonize both models (Figure A10) demonstrated significant differences in the structure of each community. For the three fecal donors studied, we observed one fecal donor (FS05) exhibited more similar community structures between ^HMA^mice and fecal inocula, one fecal donor (FS04) exhibited more similar community structure between MBRA communities and the fecal inoculum, and the third fecal donor (FS06) had community structures in MBRA and ^HMA^mice that were equally dissimilar to the starting inoculum. Examining differences in specific taxa between models also illustrated these differences. For example, in mice, *Turicibacter* was present in 55% of samples, composing ∼0.7% of reads overall. It decreased in relative abundance upon azithromycin and clindamycin treatment. In bioreactors, however, it was detected in only 0.38% samples and comprised 0.00025% of reads. Similarly, *Dialister*, which was present in 33% of MBRA communities and represented ∼0.5% of total reads was significantly lower in ceftriaxone-treated communities compared to untreated controls. However, *Dialister* was found in <0.4% of mouse samples and represented only 5.7 X 10^-5^% of reads overall. Other bacteria, like *Faecalibacterium* abundant in all of the fecal samples studied here (Huang et al., 2025), poorly colonize both models and therefore their responses to antibiotics were unable to be assessed.

### Comparison of results to commonly understood taxonomic spectrum of antibiotics

We compared our results to other studies on the effects of antibiotics on microbiota to better understand how our models compare to previously published data and known antimicrobial spectrums. Overall we observed several trends that were consistent between our models and previously published studies of effects on the GI microbiota, but we also noted differences. This was not surprising, as even studies with different populations of human subjects report different observations. Below we focus on two antibiotics, vancomycin and metronidazole, with a more complete discussion of all antibiotics studied in Appendix 1.

Vancomycin is known to affect Gram-positive cell walls and is most commonly used to target infections of the Bacillota *Staphylococcus* or *Clostridioides difficile* (Levine, 2006). Gram-negative bacteria such as the Bacteroidota are generally thought to be resistant, and low concentrations of vancomycin are even used in some nominally *Bacteroides*-selective medias. Although vancomycin has been described as narrow spectrum (Hermans and Wilhelm, 1987), previous studies (Isaac et al., 2017a; Lewis et al., 2015; Rea et al., 2011) along with the work described here demonstrates its broad effects on the microbiome. Some oral *Bacteroides* isolates are known to be sensitive to vancomycin (Van Winkelhoff and De Graaff, 1983), and there was a decreased relative abundance of Bacteroidota observed in mouse feces (Ajami et al., 2018), *C. difficile* infected mice (Lewis et al., 2015; Yamaguchi et al., 2020), and human volunteers (Isaac et al., 2017b; Russell et al., 2013) treated with vancomycin. The wide spectrum of vancomycin activity may contribute to its serious side-affects, such as increased susceptibility to developing asthma (Russell et al., 2013), risk of recurrence of *C. difficile* infection (Cornely et al., 2013), and association with increased prevalence of vancomycin-resistant enterococci (Fridkin et al., 2001).

Although Gram-positive bacteria may be more susceptible to low concentrations of vancomycin, at higher concentrations, Gram-negative bacteria may be equally affected. Vancomycin is a hydrophilic glycopeptide. At low concentrations, hydrophilic molecules primarily travel through the outer membrane via porins, however vancomycin is too large (Yarlagadda et al., 2016). Other less characterized mechanisms are responsible for cell entry at higher concentrations. In the presence of bile salts, cell membranes will have increased permeability (Begley et al., 2005) and therefore are possibly more susceptible to alternate modes of entry. This observation may explain the higher sensitivity to vancomycin treatment that we observed for control strains grown in BRM3 (which contains bovine bile) compared to YCFA (which lacks bile), although differences in other nutrients between the two media may also contribute to these observations. Metronidazole does not accumulate to high levels in feces of patients undergoing standard metronidazole treatment (1-24 mg/g stool (Bolton and Culshaw, 1986)), so the concentration used for treatment here (200 μg/ml) was higher than would be expected and effects are likely to be of reduced magnitude in patients. The partial recovery in diversity observed after cessation of antibiotic treatment we observed in MBRA communities is consistent with rapid microbiota recovery following cessation of treatment in C57Bl/6 mice (Lewis et al., 2015), although recovery following treatment in dogs (Belchik et al., 2024) and children (Gotfred-Rasmussen et al., 2021) was longer (1-2 months). We also observed similar increases in levels of *Bifidobacterium*, *Enterobacteriaceae* (*Escherichia-Shigella*), and *Enterococcus* to those reported in previous studies in dogs and rats (Marshall-Jones et al., 2024; Pélissier et al., 2010; Pilla et al., 2020).

### Potential loss of taxa secondarily affected by antibiotics

We observed several taxa with lower levels of relative abundance following treatment with many classes of antibiotics. While some of these taxa may be highly susceptible to antibiotics, other taxa may have had lower levels of relative abundance across multiple antibiotic treatments because they were reliant on commensal or synergistic microbial interactions lost during treatment. *A. muciniphila* was one taxa that was significantly lower in MBRA communities treated with ceftriaxone, ciprofloxacin, clindamycin, doxycycline, imipenem, and metronidazole but was not significantly lower in antibiotic-treated ^HMA^mice. Previous characterization of *A. muciniphila* indicated that this species is susceptible to imipenem (Dubourg et al., 2017, 2013), ceftriaxone (Dubourg et al., 2017), and doxycycline (Dubourg et al., 2013), but resistant to clindamycin (Cozzolino et al., 2020), metronidazole (Dubourg et al., 2013), and ciprofloxacin (Filardi et al., 2022). Treatment with ciprofloxacin (Rodriguez-Ruiz et al., 2024) and clindamycin (Buffie et al., 2012) also led to increases in *Akkermansia* in patients and mice, respectively, consistent with resistance to these antibiotics.

Pure culture isolates of *A. muciniphila* from human (BAA-835) and mouse (YL44) samples grew poorly in monoculture in bioreactor medium (BRM3), but grew well in the rich medium, YCFA (Figure A11). Addition of the mixture of short chain fatty acids (SCFA) and vitamins found in YCFA to BRM3 allowed for partial (*A. muciniphila* BAA-835) or full (*A. muciniphila* YL44) restoration of growth to levels observed in YCFA. This data suggests that BRM3 lacks factor(s) important for *A. muciniphila* growth that can be provided by YCFA. We propose that these factors are likely also provided by other members of the community when grown in a mixed population. Consistent with a potential role for SCFA supporting growth of *A. muciniphila* in mixed communities, we observed significantly lower levels of acetate and butyrate in clindamycin-treated communities compared to untreated MBRA communities when we measured levels of these metabolites at the end of antibiotic treatment in a subset of antibiotic-treated communities (Figure A12).

While *A. muciniphila* itself has been proposed to be a keystone species, as both *Anaerostipes caccae* (Chia et al., 2018) and *Clostridium difficile* (Engevik et al., 2020) have been demonstrated to benefit from coculture with *A. muciniphila*, there is little known about commensal interactions between *A. muciniphila* and other bacteria. One study demonstrated that the presence of *Parabacteroides merdae* leads to enhanced growth of *A. muciniphila* (Olson et al., 2018), but the nature of this interaction was not characterized.

While these data are consistent with the hypothesis that *A. muciniphila* can be lost in MBRA communities due to loss of commensal interactions with antibiotic-susceptible microbes, the alternative hypothesis that strains present in these communities were susceptible to antibiotics cannot be disproved. Although not statistically significant, *Akkermansia* levels were also lower in clindamycin-treated ^HMA^mice. Vancomycin treatment led to lower levels of many taxa that were also reduced following treatment with ceftriaxone, ciprofloxacin, clindamycin, imipenem, and metronidazole, but lower levels of *Akkermansia* was not observed. Previous studies demonstrated resistance of *A. muciniphila* to vancomycin (Dubourg et al., 2013). Future studies are needed to identify the role of potential synergistic or commensal interactions between *Akkermansia* and other gut microbes during growth in conditions that may exist in the lumen of the GI tract.

## CONCLUSIONS

In summary, this work highlights the importance of testing antibiotics on complex communities of microbes under conditions that mimic the gastrointestinal environment. Testing complex communities facilitates identification of potential context-dependent interactions that increase antibiotic susceptibility among taxa that would not be predicted based on testing with more limited panels of isolated strains. This work also demonstrates the strengths and weaknesses of the two models tested. Both models reproduced some of the observed effects of antibiotics on the microbiota reported in human and other animal studies. However, key differences were also observed. While this is not surprising, as many of the studies cited above also had conflicting observations, it still limits the ability to drawn firm conclusions about the effects of each antibiotic on the microbiota. Both models were also limited in their ability to fully recapitulate the complexity of communities found in human fecal samples. While minibioreactors allowed more fecal samples to be tested in higher throughput at reduced cost, effects of antibiotics that were influenced by pharmacokinetics, host metabolites, and/or microbiome-immune interactions could not be studied. While ^HMA^mice provide these more complex interactions, more extensive experimentation was limited by ethical concerns to reduce the number of animals used in research (Hubrecht and Carter, 2019) and the higher costs associated with animal experimentation. Additionally, the compositional variation introduced as a result of interactions between mouse genetic background and microbiota composition indicated the potential importance of strain background in understanding the effects of antibiotics on the microbiota when working with ^HMA^mice. Nevertheless, these data demonstrate that both models can be useful pre-clinical screening tools during the characterization of new antimicrobial compounds to combat the threats associated with continued spread of antibiotic resistance (Talbot et al., 2006).

## DATA AVAILABILITY

16S rRNA gene data has been deposited in NCBI’s Sequence Read Archive (SRA) under BioProject ID PRJNA729569. Code for analysis of data is available at Zenodo: https://zenodo.org/records/14728946; doi:10.5281/zenodo.14728946.

## AUTHOR CONTRIBUTIONS

J.M.A., T.AA., and A.I.L. were responsible for conceptualization and design of the studies. Data collection was performed by T.A.A., A.I.L, and K.S. Analysis was performed by T.A.A, A.I.L, and J.M.A. J.M.A. acquired funding for the studies. The initial draft of the manuscript was written by T.A.A. and J.M.A., and all authors revised, edited, and approved the final manuscript.

## ACKNOWLEDGEMENTS

We thank Hugh McCullough, Austin Johnson, Yining He, and Delphin Mutangana for technical assistance with MBRA experiments and growth of *A. muchiniphila* strains. We thank Amanda Ramer-Tait, Robert Schmaltz, and Rafael Segura-Muñoz of the Nebraska Gnotobiotic mouse program and Kylie Farrell, formerly of Baylor College of Medicine, for their assistance in establishing ^HMA^mice models. We thank Devin Rose (University of Nebraska-Lincoln) for use of HPLC for analysis of SCFA. We also acknowledge and Kristin Beede and Paula Chacua for providing or assisting in isolation of strains. Sequence analysis was completed utilizing the Holland Computing Center, which receives support from the UNL Office of Research and Economic Development and the Nebraska Research Initiative. This research was supported by CDC contracts 200-2017-96080 and 75D301-18C-02909, as well as Nebraska Tobacco Settlement Biomedical Research Development Funds awarded to J.M.A.

## CONFLICTS OF INTEREST

J.M.A. and T.A.A. have a significant financial interest in Synbiotic Health. There are no other conflicts of interest to declare.

## ETHICS APPROVAL

Protocols for collection and use of fecal samples were reviewed and approved by Institutional Review Boards at Baylor College of Medicine (protocol number H-38014) and University of Nebraska-Lincoln (protocol number 18585). Experiments with mice were performed according to protocols approved by the Institutional Animal Care and Use Committees of Baylor College of Medicine (AN-6675) and University of Nebraska-Lincoln (IACUC #1668, #1680) and are in accordance with the NIH Office of Laboratory and Animal Welfare (OLAW) guidelines.

## Appendix

**Figure A1.**
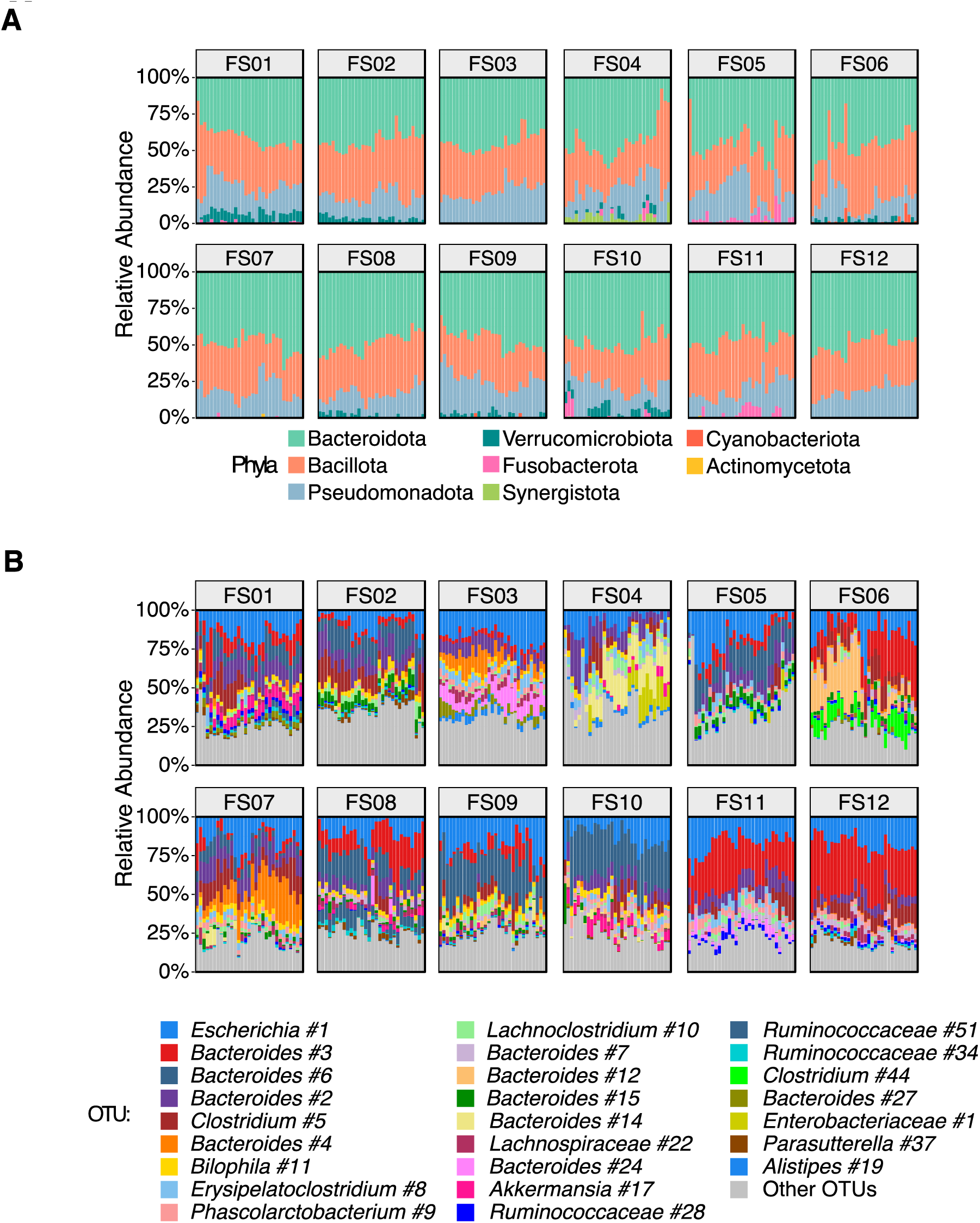
Relative abundance of taxa in MBRA communities prior to treatment. Relative abundance of (A) phyla and (B) twenty-five most abundant OTUs across replicate MBRA communities prior to treatment on Day 0.

**Figure A2:**
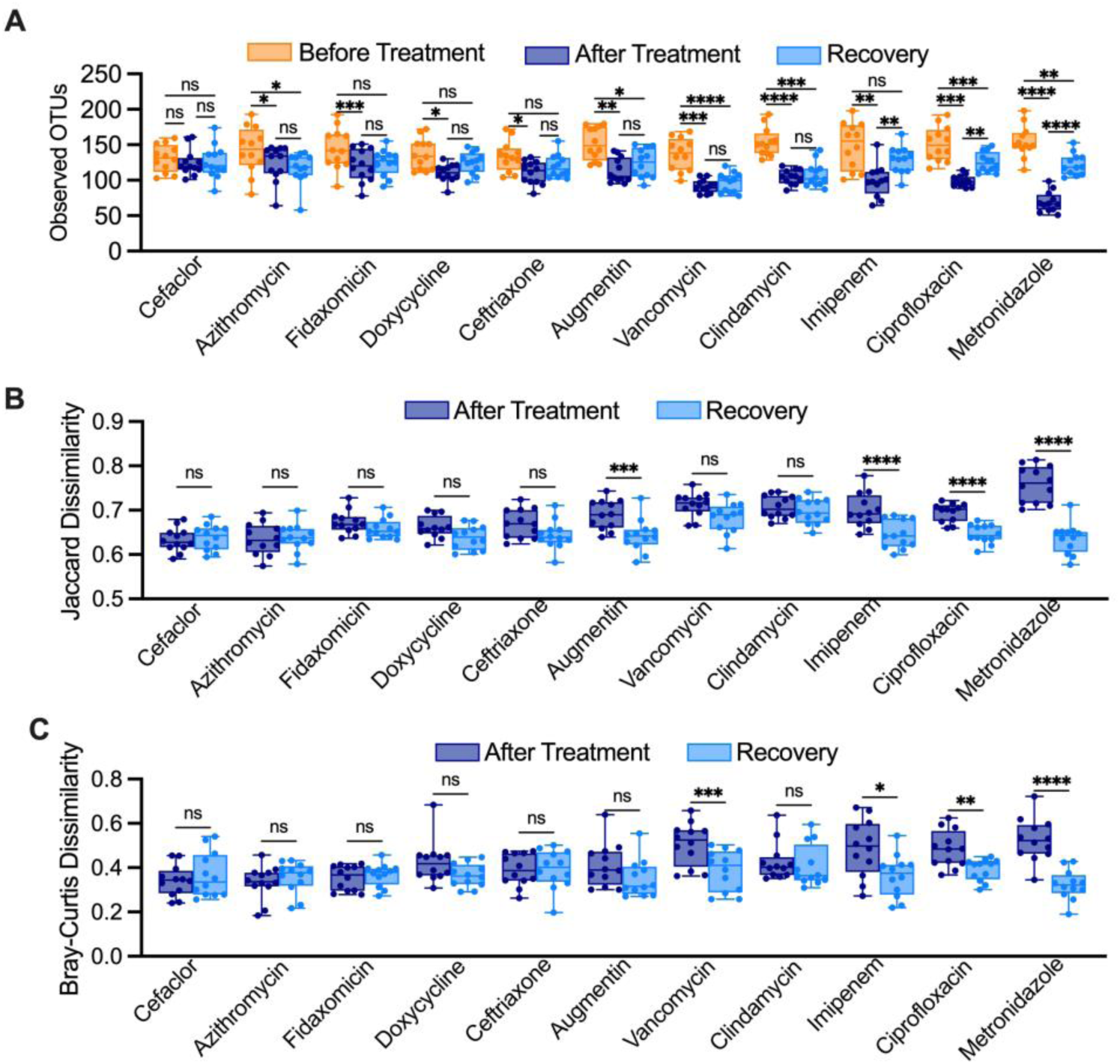
Partial recovery of microbial diversity following treatment with a subset of antibiotics for human fecal communities cultured in MBRAs. (A) Observed OTUs assessed two days following the end of treatment (Recovery, Day 7) were plotted along with data collected before (Day 0) and after (Day 5) treatment with the treatment indicated on the x-axis. Individual points were plotted from the mean of replicates for each of the twelve fecal samples tested. (B) Jaccard and (C) Bray-Curtis dissimilarity values from samples before treatment (Day 0) were determined for samples collected after treatment (Day 0 -Day 5) or after a two day recovery following antibiotic treatment (Day 0 - Day 7), with the treatment indicated on the x-axis. Individual points were plotted from the mean of replicate dissimilarity measures for each of the twelve fecal samples tested. In all panels, data from before and after treatment were reproduced from Figure 1, with the exception of data from cefaclor and ceftriaxone-treated communities, which were restricted to duplicate samples collected from reactors where community composition was measured on Day 7. Median values were plotted as horizontal lines, interquartile (IQR) ranges were plotted as boxes, and range of data was plotted as whiskers. Statistical significance was determined by paired ANOVA with Geisser-Greenhouse correction for unequal variability of differences and Holm-Šidák correction for multiple comparisons. In (A), comparisons were performed between all time points from the same treatment. In (B) and (C), comparisons were made between dissimilarity values for the same treatment. ns>0.05; *, p<0.05; **, p<0.01; ***, p<0.001; ****, p<0.0001.

**Figure A3.**
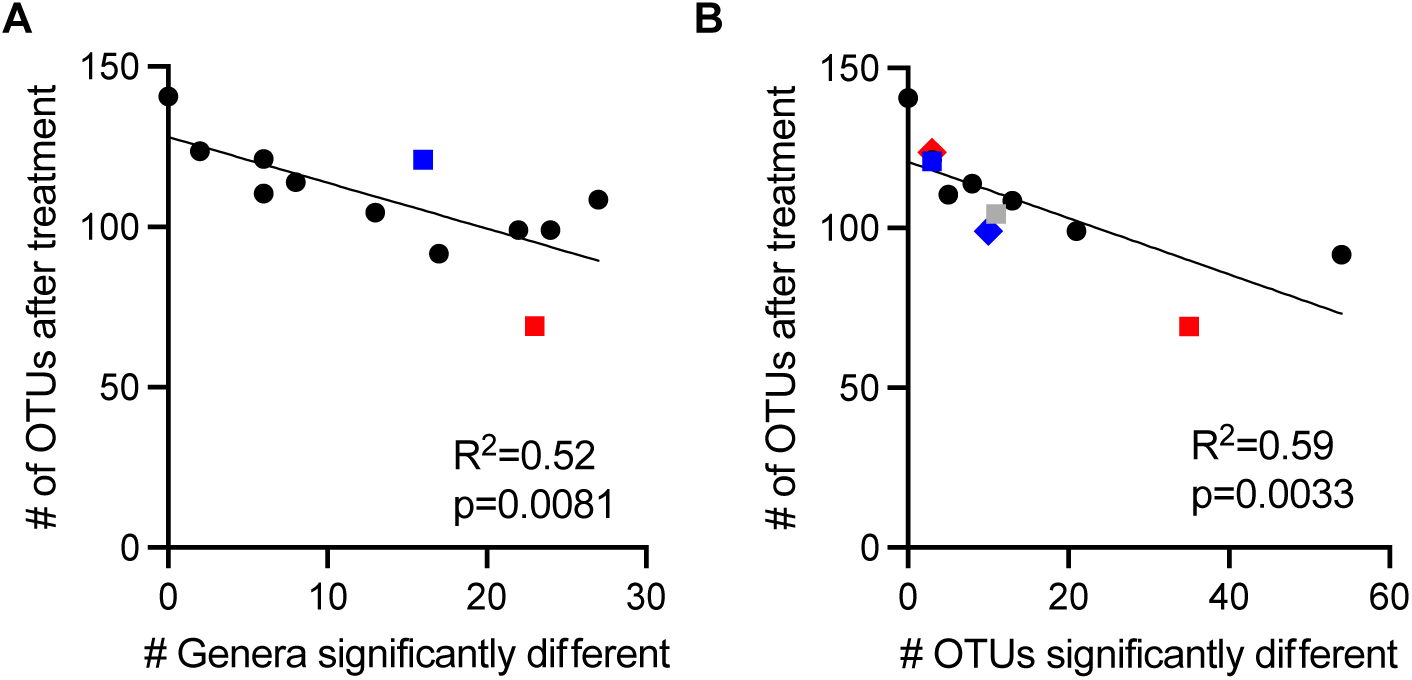
Number and diversity of taxa loss were correlated. We plotted the number of OTUs remaining after treatment with an antibiotic as a function of the number of (A) genera or (B) OTUs that were significantly different between communities treated with that antibiotic and untreated communities. Treatments that were above or below the 95% confidence interval of the correlation are indicated: fidaxomicin, blue square; metronidazole, red square; azithromycin, red diamond; ciprofloxacin, blue diamond; clindamycin, gray square; cefaclor, gray diamond (obscured by fidaxomicin and azithromycin).

**Figure A4.**
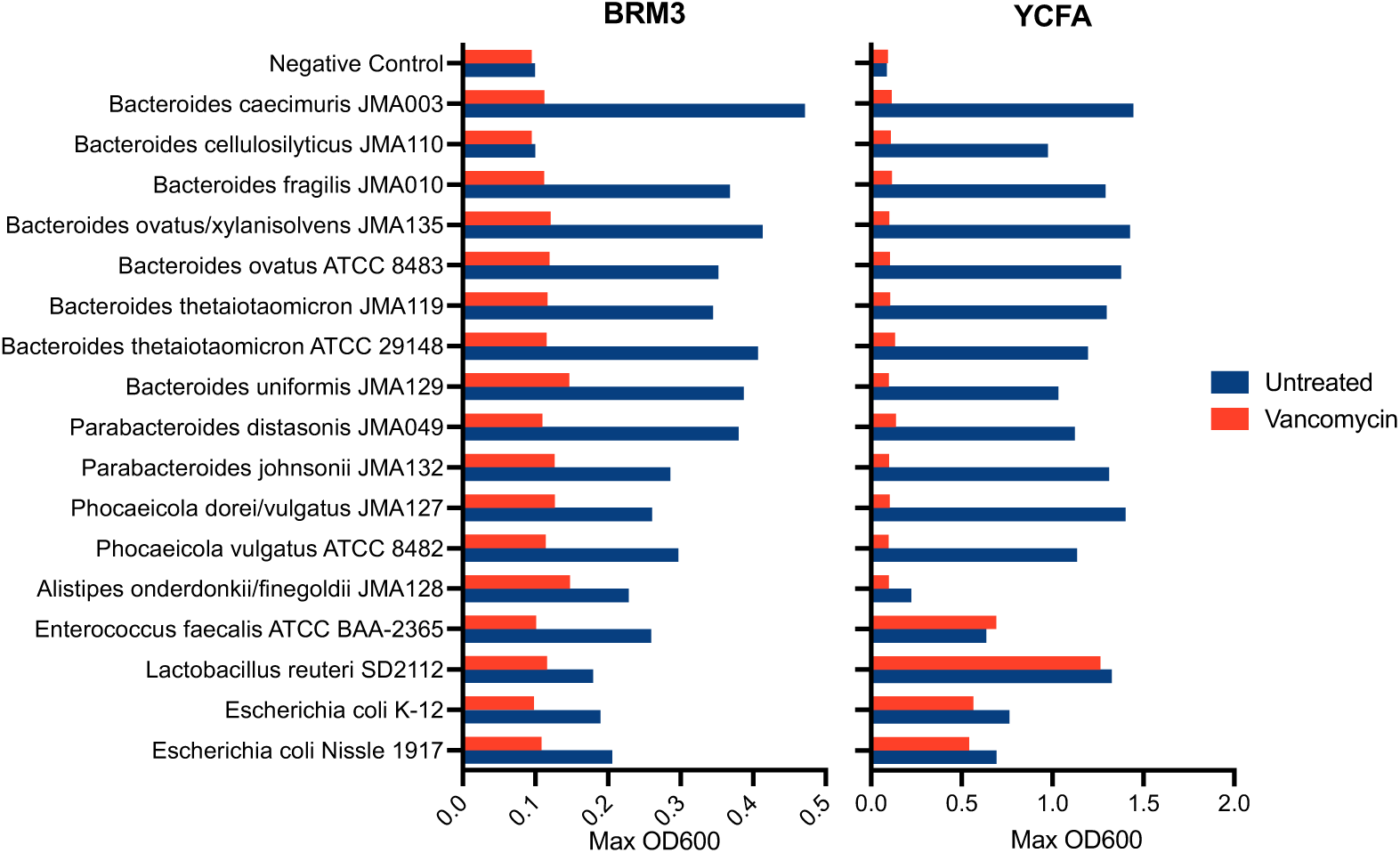
Growth of Bacteroidota isolates and control strains in medias with and without vancomycin. Displayed are the maximum optical density at 600 nm of 48 hr growth curves in BRM3 or YCFA, with (red) or without (blue) 150 μg/ml vancomycin. Data points are an average of two replicates. *Limosilactobacillus reuteri* was grown on MRS instead of YCFA. The MRS negative controls had a maximum OD of 0.13 (untreated). Isolates are indicated by the strain name “JMA” followed by a number. Taxonomic names for JMA strains were based on 16S rRNA gene sequencing as described in Methods.

**Figure A5.**
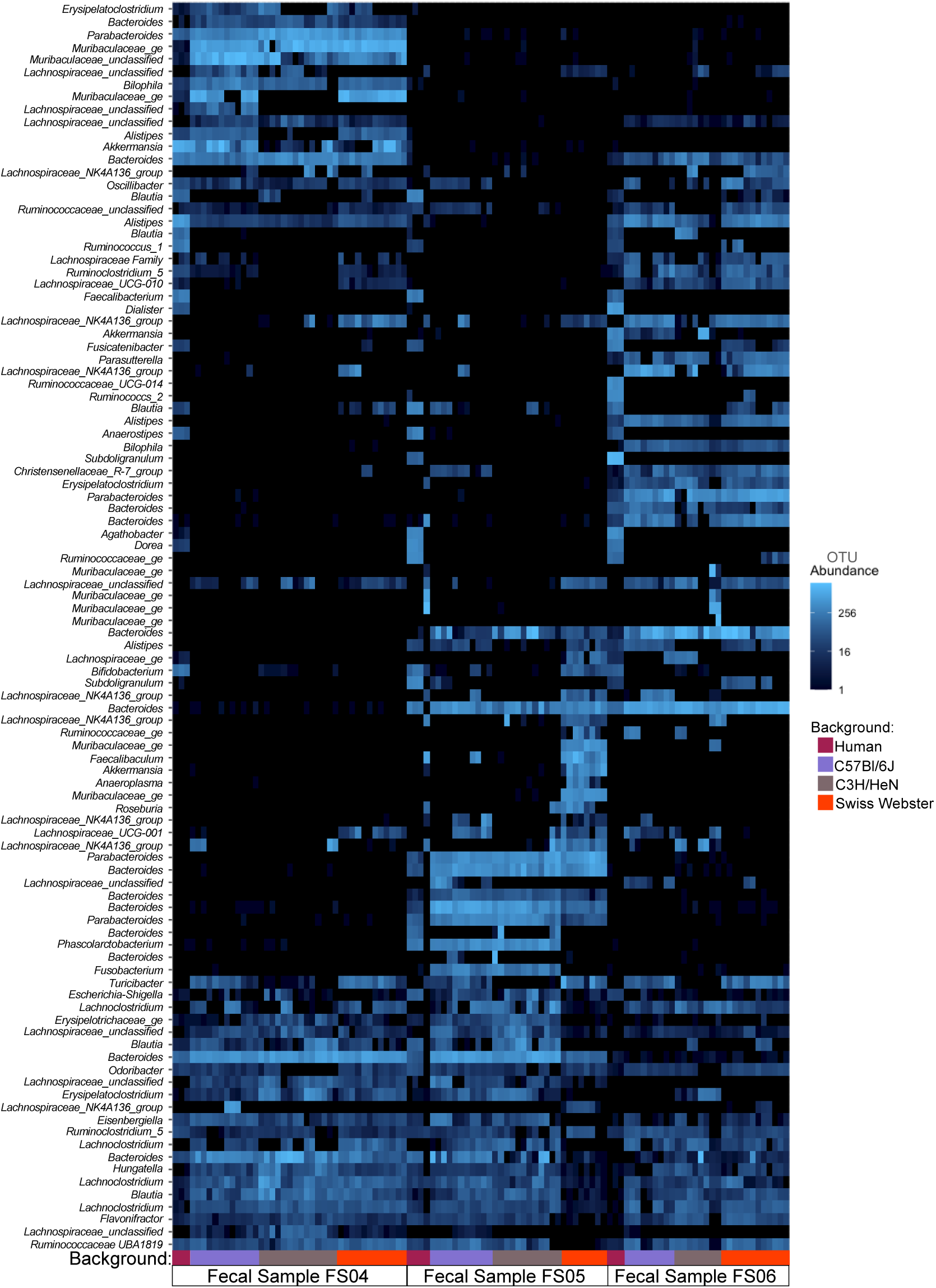
100 most abundant OTUs across human fecal samples used for colonization and collected from untreated HMAmice colonized with indicated fecal samples. Fecal sample and mouse genetic background are indicated on the x-axis. The y-axis indicates genus-level classification of OTU (or lowest taxonomic rank assigned with confidence). Abundance varies based on shading as indicated.

**Figure A6.**
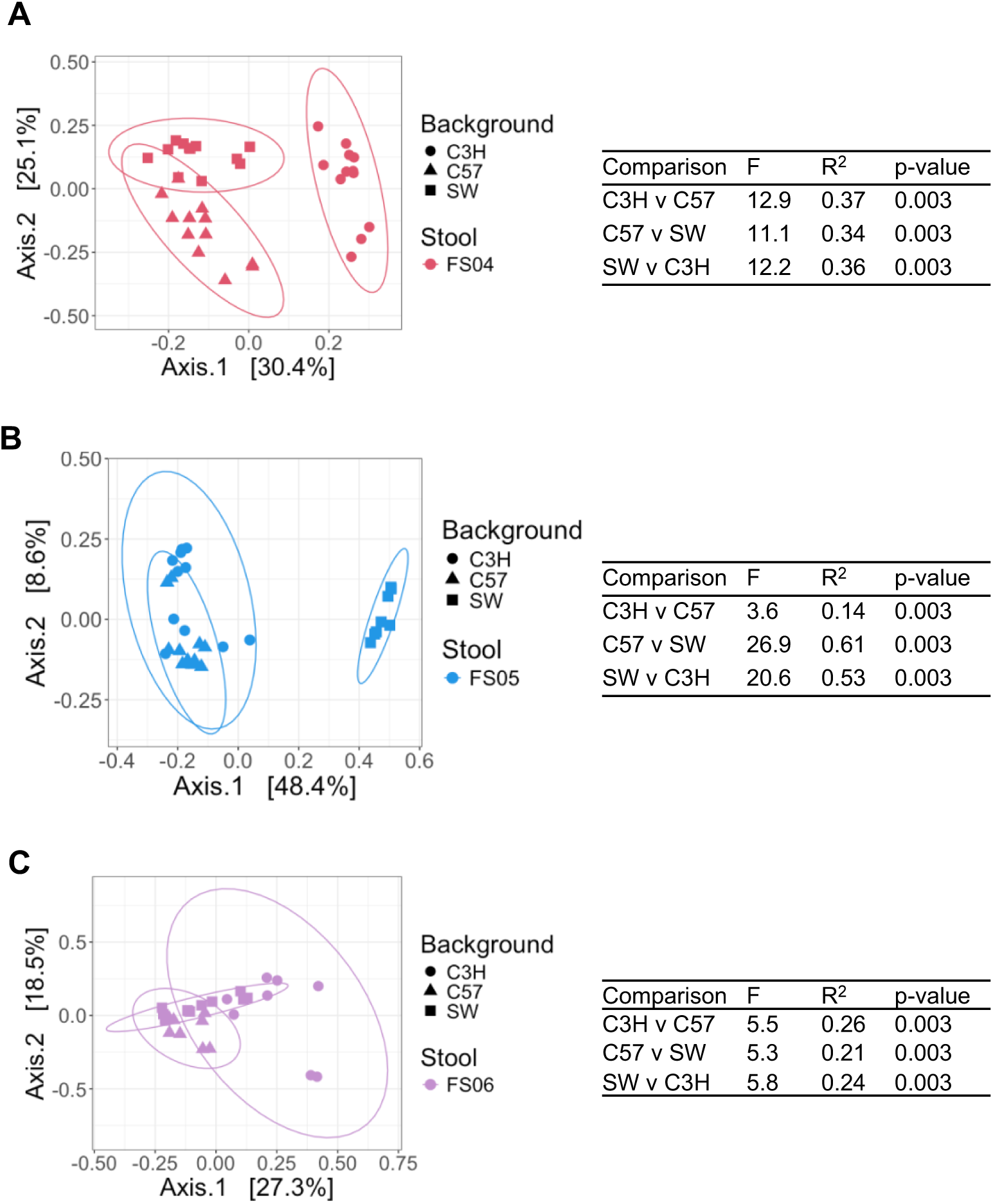
Microbiota structure varies by mouse genetic background in untreated mouse fecal communities. PCoA plots of Bray-Curtis dissimilarities for samples collected from untreated mice colonized with (A) FS04, (B) FS05, or (C) FS06. Shapes indicated mouse genetic background. Statistical significance of differences between groups was determined with pairwise ANOVA, with statistic indicated to the right of each panel. Ellipses indicate 95% confidence intervals.

**Figure A7.**
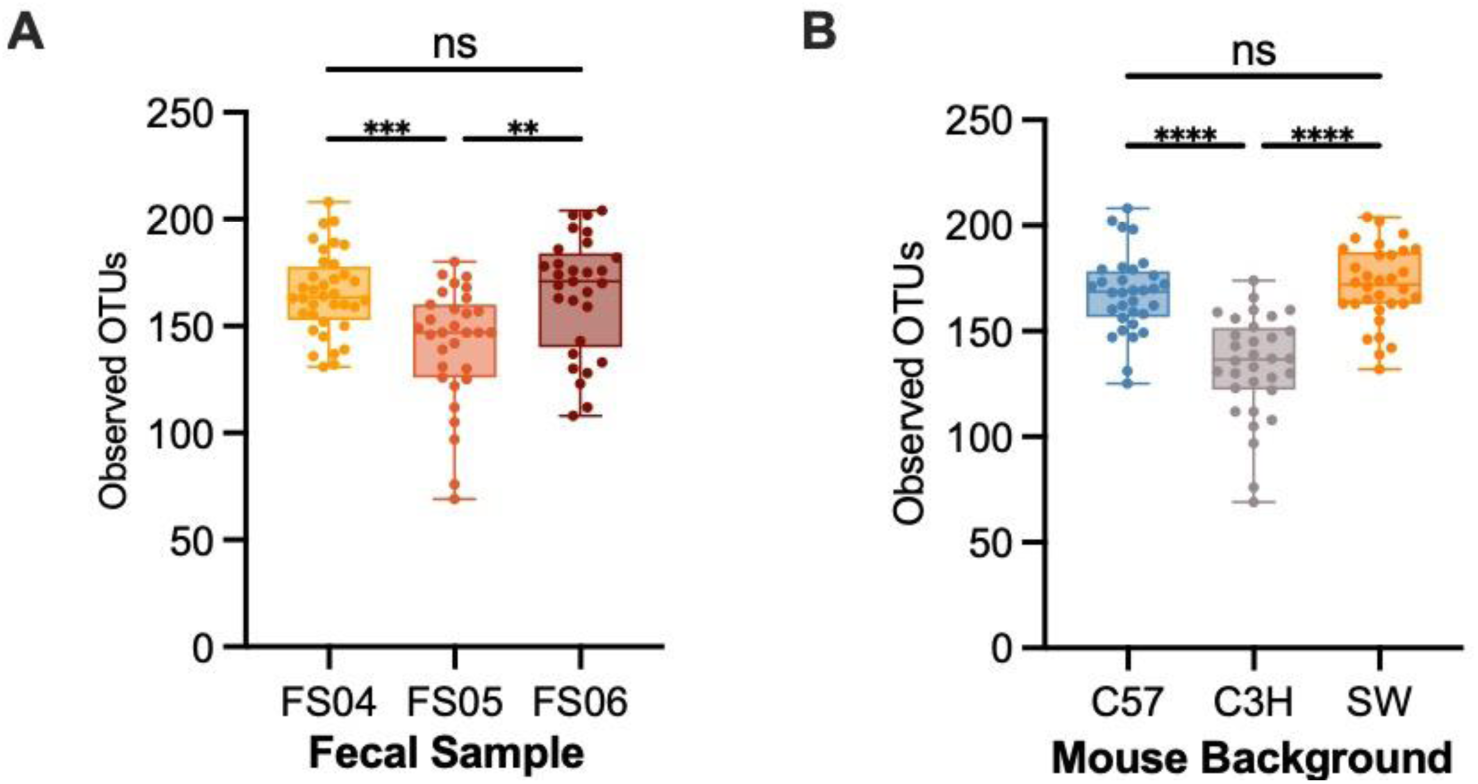
Levels of observed OTUs vary in the fecal samples of ^HMA^mice based upon both fecal donor and mouse genetic background. As in Figure 4, observed OTUs were plotted from mice in the untreated group after five days of mock-treatment with PBS. Levels of OTU abundance based on (A) fecal donor used for colonization or (B) mouse genetic background were plotted. Statisical significance of differences between groups was determined with one-way ANOVA with Brown-Forsythe and Welch correction for unequal variances and Dunnett’s T3 correction for multiple comparisons.

**Figure A8.**
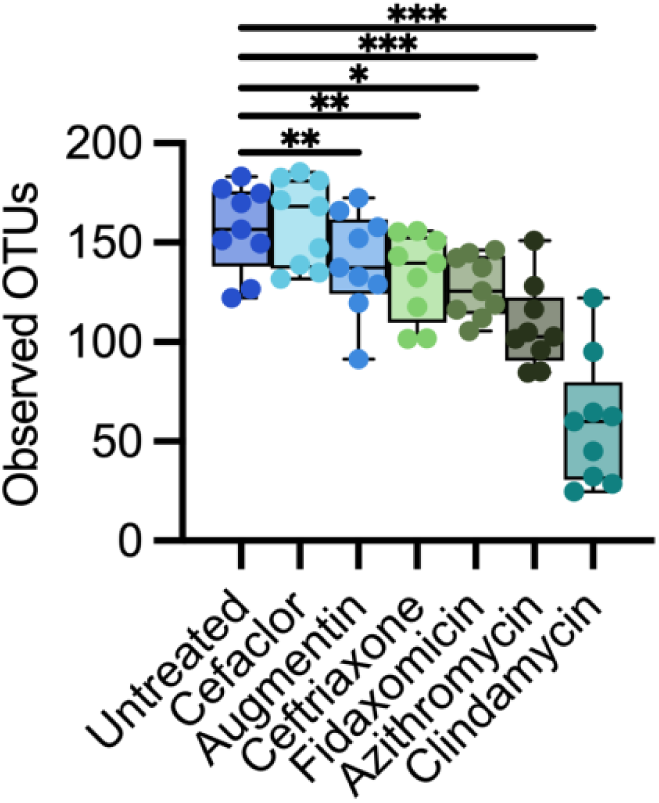
Changes in Observed OTUs across all antibiotic treated mice compared to controls. Mean levels of observed OTUs for each mouse genetic background/fecal donor (n=9) were plotted by treatment group. Statistical significance of differences in levels observed compared to untreated were calculated was determined by paired ANOVA with Geisser-Greenhouse correction for unequal variability of differences and Holm-Šidák correction for multiple comparisons. *, p<0.05; **, p<0.01; ***, p<0.001; ****, p<0.0001.

**Figure A9.**
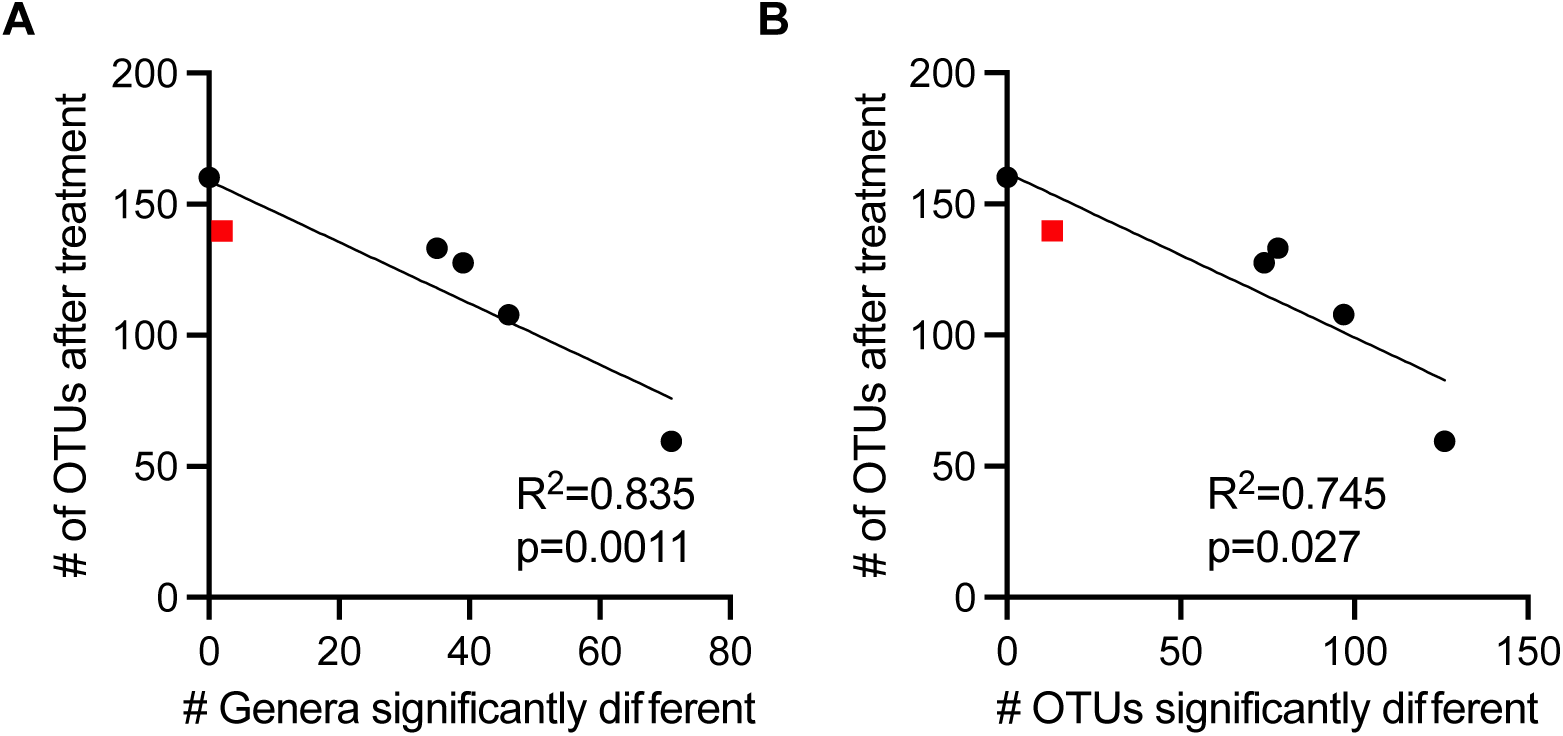
Number and diversity of taxa loss were correlated in ^HMA^mice. We plotted the number of OTUs remaining after treatment with an antibiotic as a function of the number of (A) genera or (B) OTUs that were significantly different between communities treated with that antibiotic and untreated communities. In both plots, only Augmentin-treated communities (red squares) were outside of the 95% confidence interval of the correlation.

**Figure A10.**
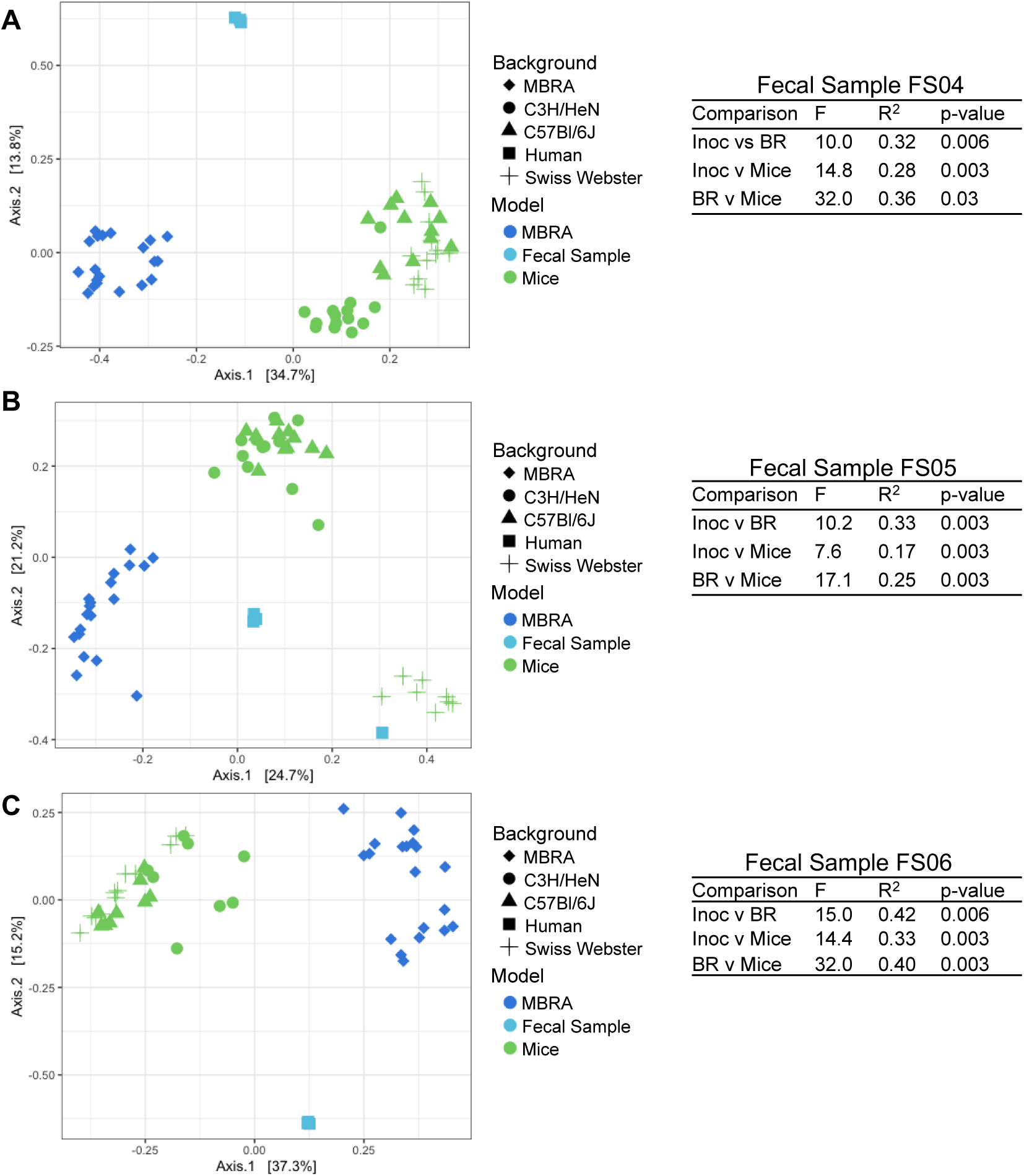
Microbiota composition in MBRA and ^HMA^mouse models of the human GI microbiota. PCoA plots of Bray-Curtis dissimilarities show differences in community structure measured between original fecal inocula (light blue squares), bioreactor samples collected before treatment (dark blue circles) and ^HMA^mice (green symbols) in the C57Bl/6J (diamonds), C3H/HeN (circles), or Swiss-Webster (triangles) mouse backgrounds for fecal sample (A) FS04, (B), FS05, or (C) FS06. Percent variation explained by each axis is indicated. Pairwise Permanova testing indicated statistically significant differences between models for each fecal sample. Abbreviations in Permanova tables are Inoc: fecal inoculum, BR: MBRA communities, Mice: ^HMA^mice.

**Figure A11.**
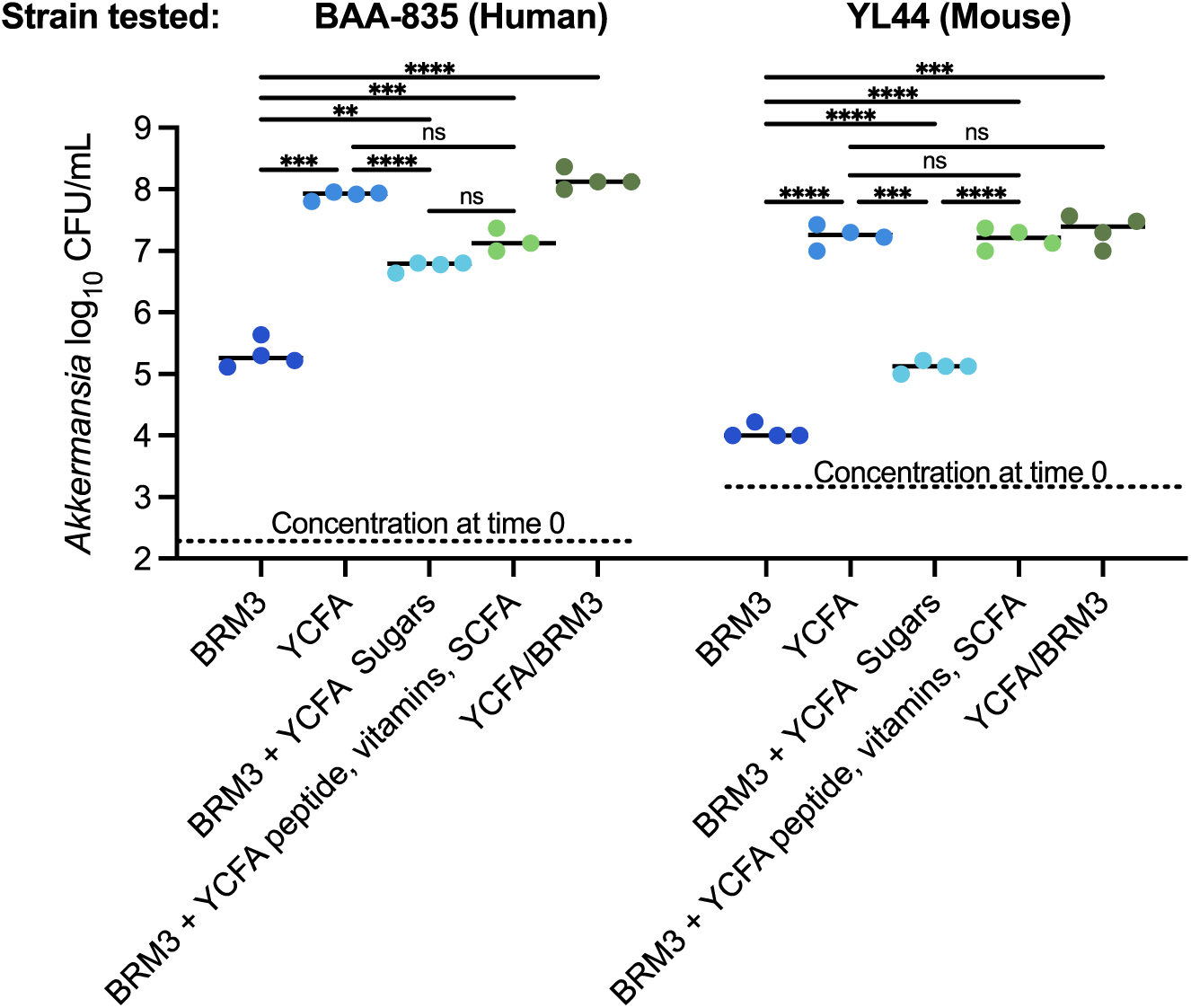
Akkermansia muciniphila requires compounds absent in BRM3 for high levels of growth. We cultured human and mouse isolates of A. muciniphila in BRM3, YCFA, and mixtures of BRM3 amended with components from YCFA as described in Table A4. Concentration of cells in the inoculum and after 24 hours of growth was determined by serial dilution and plating as described. Statistical significance of differences between growth media for each strain was determined by one-way ANOVA with Brown-Forsythe and Welch correction for unequal variances and Dunnett’s T3 correction for multiple comparisons. ns, p>0.05; **, p<0.01; ***, p<0.001; ****, p<0.0001.

**Figure A12.**
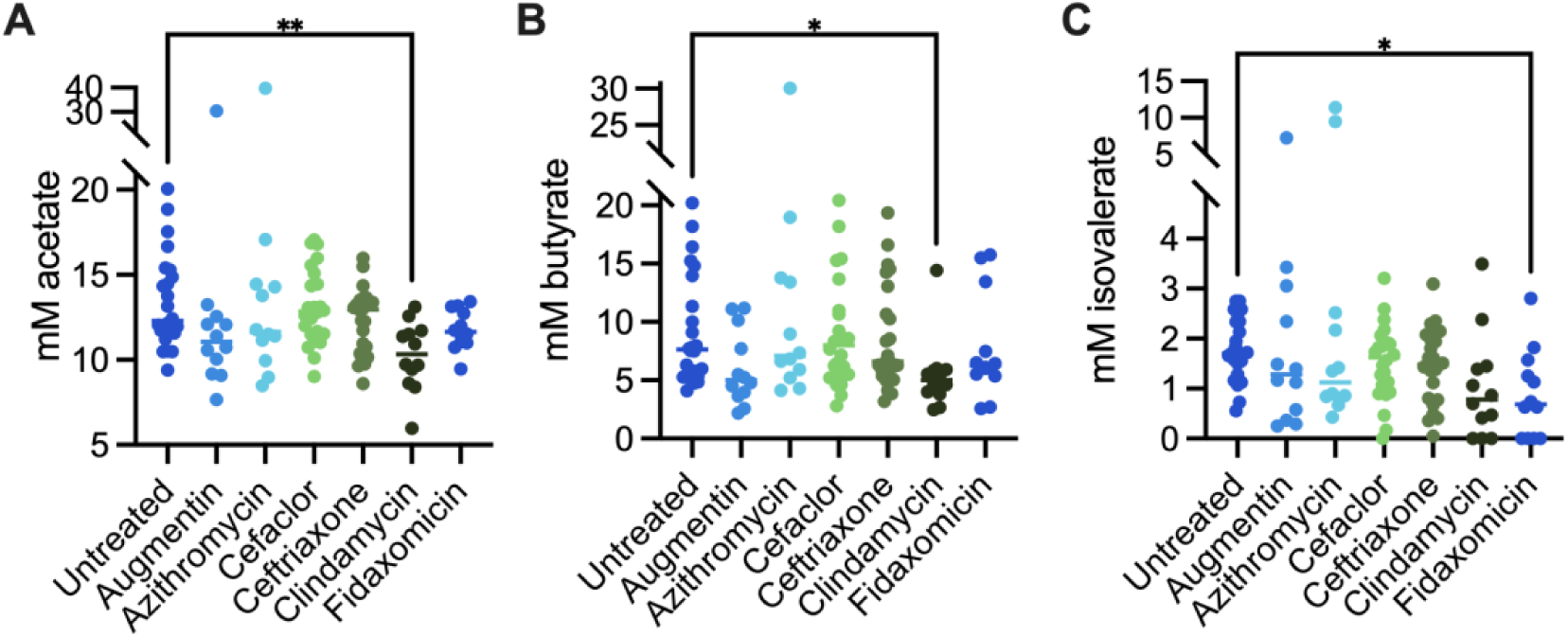
Levels of acetate and butyrate were lower in clindamycin-treated communities. We measured levels of (A) acetate, (B) butyrate and (C) isovalerate in filter-sterilized spent culture medium from MBRA communities on Day 5 in untreated (n=24) and antibiotic-treated communities (n=12, except cefaclor and ceftriaxone, where n=24). Statistical significance of differences between antibiotic-treated and untreated communities was determined with one-way ANOVA with Brown-Forsythe and Welch correction for unequal variances and Dunnett’s correction for multiple comparisons. *, p<0.05; **, p<0.01. Comparisons to untreated with p>0.05 are not shown.

**Table A1:** Sources of antibiotics used in this study.

|  |  |
| --- | --- |
| Augmentin<br>(5:1 amoxicillin sodium salt to potassium clavulanate) | Research Products International |
| Azithromycin dihydrate | Thermo Scientific Chemicals |
| Cefaclor | Thermo Scientific Chemicals |
| Ceftriaxone sodium salt hemiheptahydrate | Thermo Scientific Chemicals |
| Clindamycin phosphate, 97+% | TCI America |
| Fidaxomicin | Apexbio Technology |
| Ciprofloxacin hydrochloride hydrate, 98% | Thermo Scientific Chemicals |
| Doxycycline monohydrate | Thermo Scientific Chemicals |
| Imipenem monohydrate, 98% | Thermo Scientific Chemicals |
| Metronidazole, 99% | Thermo Scientific Chemicals |
| Sulfamethoxazole, 98+% | TCI America |
| Vancomycin hydrochloride, USP grade | VWR |

**Table A2:** Number of mice per treatment group*.

|  | **C57Bl6/J** |  |  | **C3H/HeN** |  |  | **Swiss Webster** |  |  |
| --- | --- | --- | --- | --- | --- | --- | --- | --- | --- |
| **Treatment:** | FS06 | FS05 | FS04 | FS06 | FS05 | FS04 | FS06 | FS05 | FS04 |
| Augmentin | 6,<br>(33%) | 6,<br>(50%) | 6,<br>(67%) | 6,<br>(50%) | 6,<br>(33%) | 6,<br>(33%) | 6,<br>(50%) | 2,<br>(50%) | 6,<br>(50%) |
| Azithromycin | 6,<br>(50%) | 6,<br>(67%) | 6,<br>(50%) | 6,<br>(67%) | 6,<br>(33%) | 6,<br>(33%) | 6,<br>(67%) | 6,<br>(33%) | 3,<br>(0%) |
| Cefaclor | 12,<br>(42%) | 11,<br>(55%) | 12,<br>(50%) | 12,<br>(67%) | 10,<br>(70%) | 12,<br>(50%) | 12,<br>(50%) | 9,<br>(33%) | 12,<br>(42%) |
| Ceftriaxone | 10,<br>(50%) | 11,<br>(27%) | 11,<br>(36%) | 12,<br>(50%) | 12,<br>(42%) | 12,<br>(58%) | 12,<br>(58%) | 9,<br>(33%) | 10,<br>(50%) |
| Clindamycin | 5,<br>(40%) | 5,<br>(60%) | 6,<br>(33%) | 5,<br>(20%) | 6,<br>(17%) | 5,<br>(80%) | 6,<br>(50%) | 6,<br>(50%) | 6,<br>(33%) |
| Fidaxomicin | 6,<br>(33%) | 6,<br>(67%) | 6,<br>(50%) | 6,<br>(67%) | 6,<br>(83%) | 6,<br>(50%) | 6,<br>(67%) | 6,<br>(50%) | 6,<br>(50%) |
| Untreated | 9,<br>(78%) | 11,<br>(73%) | 12,<br>(50%) | 8,<br>(100%) | 12,<br>(42%) | 12,<br>(33%) | 12,<br>(42%) | 8,<br>(38%) | 12,<br>(33%) |

\*Percent of female mice indicated in parentheses.

**Table A3:**
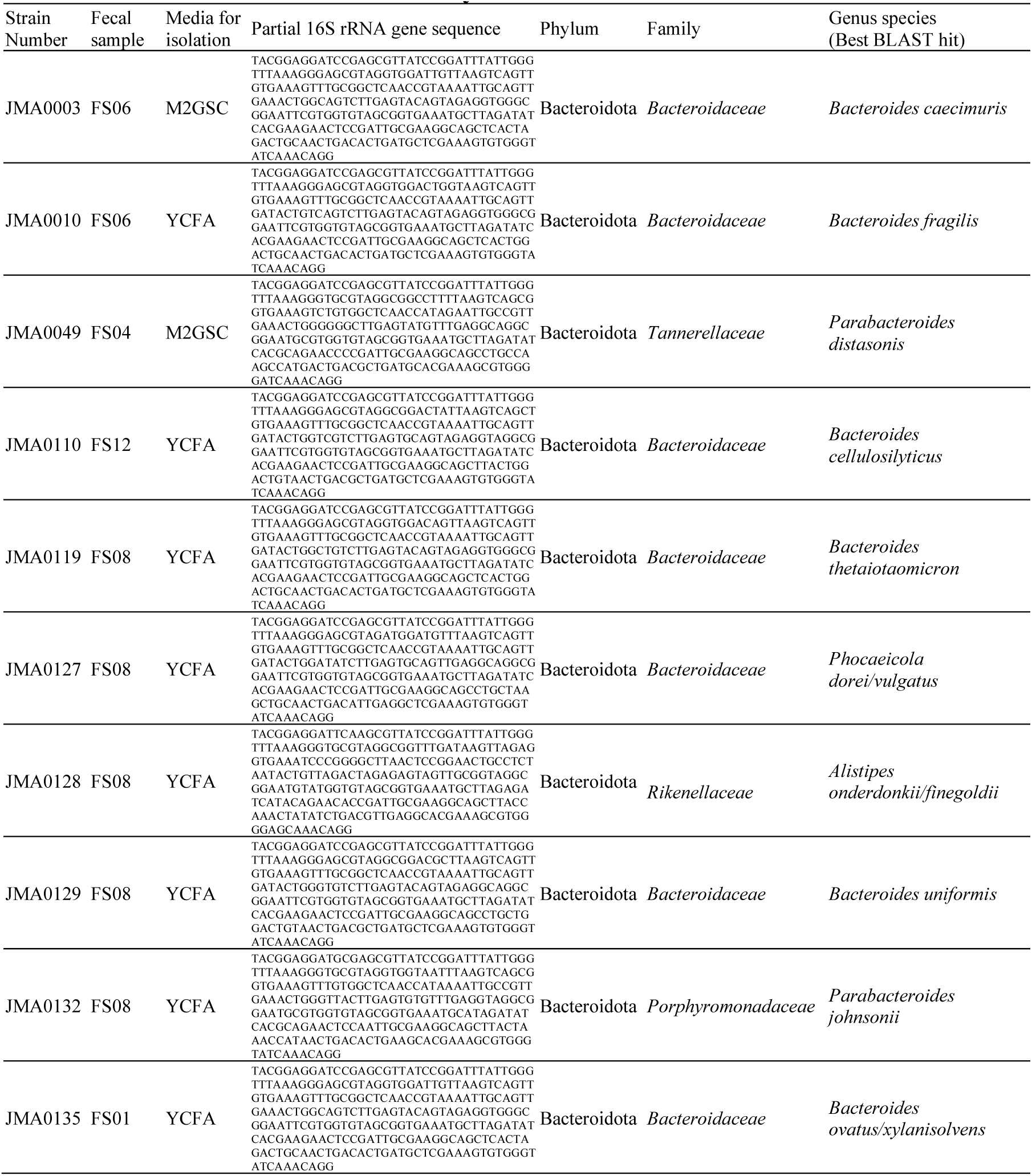
Strains isolated in this study.

**Table A4.** Variations of media used for *Akkermansia muciniphila* growth.

|  | YCFA | BRM3 | BRM3+YCFA<br>Sugars | BRM3+YCFA<br>peptide,<br>vitamins, and<br>SCFA | BRM3+YCFA |
| --- | --- | --- | --- | --- | --- |
| casitone | 10 g | - | - | 10 g | 10 g |
| tryptone | - | 1 g | 1 g | 1 g | 1 g |
| proteose peptone #3 |  | 2 g | 2 g | 2 g | 2 g |
| yeast extract | 2.5 g | 2 g | 2 g | 2.5 g | 2.5 g |
| cysteine | 1 g | - | - | 1 g | 1 g |
| sodium chloride | 0.9 g | 0.4 g | 0.4 g | 0.9 g | 0.9 g |
| magnesium sulfate | 0.09 g | 0.01 g | 0.01 g | 0.09 g | 0.09 g |
| calcium chloride | 0.09 g | 0.01 g | 0.01 g | 0.09 g | 0.09 g |
| bovine bile | - | 0.5 g | 0.5 g | 0.5 g | 0.5 g |
| Tween 80 | - | 2 ml | 2 ml | 2 ml | 2 ml |
| biotin | 10 µg | - | - | 10 µg | 10 µg |
| cobalamin | 10 µg | - | - | 10 µg | 10 µg |
| p-aminobenzoic acid | 30 µg | - | - | 30 µg | 30 µg |
| folic acid | 50 µg | - | - | 50 µg | 50 µg |
| pyridoxamine | 150 µg | - | - | 150 µg | 150 ug |
| vitamin K1 | 0.5 mg | 0.1 mg | 0.1 mg | 0.5 mg | 0.5 mg |
| hemin | 5 mg | 5 mg | 5 mg | 5 mg | 5 mg |
| thiamine | 2 µg | - | - | 2 µg | 2 µg |
| riboflavin | 2 µg | - | - | 2 µg | 2 µg |
| sodium acetate | 2.7 g | - | - | 2.7 g | 2.7 g |
| sodium propionate | 0.86 g | - | - | 0.86 g | 0.86 g |
| isobutyrate | 1 mM | - | - | 1 mM | 1 mM |
| isovalerate | 1 mM | - | - | 1 mM | 1 mM |
| valerate | 1 mM | - | - | 1 mM | 1 mM |
| sodium bicarbonate | 4 g | 2 g | 2 g | 4 g | 4 g |
| K<sub>2</sub>HPO<sub>4</sub> | 0.45 g | 0.04 g | 0.04 g | 0.45 g | 0.45 g |
| KH<sub>2</sub>PO<sub>4</sub> | 0.45 g | 0.04 g | 0.04 g | 0.45 g | 0.45 g |
| glucose | 2 g | 0.04 g | 2 g | 0.04 g | 2 g |
| maltose | 2 g | 0.15 g | 2 g | 0.15 g | 2 g |
| cellobiose | 2 g | 0.15 g | 2 g | 0.15 g | 2 g |
| arabinogalactan | - | 0.1 g | 0.1 g | 0.1 g | 0.1 g |
| inulin | - | 0.2 g | 0.2 g | 0.2 g | 0.2 g |
| pH | 6.8 | 6.8 | 6.8 | 6.8 | 6.8 |

**Table A5:** Pairwise ANOVA results comparing fecal donor and background in ^HMA^mice.

| *Comparisons by fecal donor* |  |  |  |
| --- | --- | --- | --- |
|  | FS05 | FS06 |  |
| FS04 | F=42.4, p=0.003 | F=52.8, p=0.003 |  |
| FS05 | NA | F=29.4, p=0.003 |  |
| *Comparisons by genetic background* |  |  |  |
|  | Human | C57Bl/6J | C3H/HeN |
| C57Bl/6J | F=8.3, p=0.006 | NA | F=3.8, p=0.024 |
| C3H/HeN | F=9.3, p=0.006 | F=3.8, p=0.024 | NA |
| Swiss Webster | F=7.7, p=0.006 | Not significant (F=3.2, p=0.12) | F=6.7, p=0.006 |

### Appendix 1: Comparisons of effects of antibiotics in GI models to previously published observations

Augmentin, a broad spectrum antibiotic that combines amoxicillin and clavulanate, is used to treat Gram-negative and Gram-positive infections (Gerber et al., 2017). A previous study in healthy adults observed decreases in *Lachnospiraceae* and *Coriobacteriaceae* and increases in *Enterobacteriaceae*, *Bacteroidaceae*, and *Porphyromonadaceae*, along with a rapid recovery of the microbiota to baseline levels one week following treatment (MacPherson et al., 2018). We observed lower levels of a *Lachnospiraceae* genus in Augmentin-treated ^HMA^mice, but did not observe any of the other changes in microbial taxa reported by MacPherson and colleagues. In the MBRA model, *Ruminococcaceae* species were significantly lower in Augmentin-treated communities relative to untreated controls, although we observed a similar return to baseline microbial community composition levels (Jaccard diversity) as was reported by MacPherson and colleagues.

Azithromycin is a broad-spectrum antibiotic with extended activity towards Gram-negative pathogens (Retsema et al., 1987). Several studies in human subjects observed loss of *Bifidobacterium* species following treatment with azithromycin (Chopyk et al., 2023; Sanyang et al., 2024; Wei et al., 2018). Parker and colleagues observed reduced levels of Pseudomonadota (Proteobacteria) and *Akkermansia* (Parker et al., 2017). Anthony and colleagues observed increases in the Bacillota genera *Ruminococcus*, *Eubacterium* and *Anaerostipes* (Anthony et al., 2022). In most studies, microbiota disruption was short-term (Anthony et al., 2022; Chopyk et al., 2023; Doan et al., 2019; Parker et al., 2017; Sanyang et al., 2024; Wei et al., 2018), although both Korpela (Korpela et al., 2016) and Anthony (Anthony et al., 2022) observed longer term disruptions. Across the models studied, azithromycin-treatment had much larger impacts on ^HMA^mice than in the MBRA model. Several responses in ^HMA^mice were consistent with observations in human subjects, including loss of *Bifidobacterium* and decreases in Pseudomonadota. Differences included loss of members of the Bacillota in ^HMA^mice and increases in levels of *Akkermansia*. The one response conserved across models was lower levels of the Pseudomonadota, *Parasutterella* in azithromycin-treated ^HMA^mice and MBRA communities. Loss of *Parasutterella* is consistent with the extended activity of this antibiotic against Gram-negative bacteria. Similar losses of *Parasutterella* have also been observed in mice treated with azithromycin (Li et al., 2017).

Cefaclor, a second generation cephalosporin, is a broad spectrum antibiotic primarily used against Gram-positive aerobes that has limited spectrum against Gram-negative pathogens (Neu and Fu, 1978). Very few GI microbiota studies have been performed in humans. In a study reported by Bennett and colleagues, two infants exhibited low levels of *E. coli*, *Bacteroides*, lactobacilli, and *Bifidobacterium* isolates (Bennet et al., 2002). Cefaclor had little (MBRA) or no (^HMA^mice) significant effects on bacterial richness or taxonomic richness.

Ceftriaxone, a third generation cephalosporin, has extended spectrum compared to cefaclor, with effects on aerobic Gram-positive and Gram-negative bacteria and limited effects on anaerobic bacteria (Richards et al., 1984). Zhao and colleagues observed decreased *Lactobacillus* and increased *E. coli* in BALB/c mice (Zhao et al., 2020). Similarly, Gregoire and colleagues observed decreased *Lactobacillus* and increased *Enterococcus* in Swiss Webster mice (Grégoire et al., 2021). Increases in *Enterococcus* were also observed in C57Bl/6J mice (de Moura e Dias et al., 2023) and Sprague-Dawley rats (Duclot et al., 2024). *Akkermansia* levels also increased in C57Bl/6J mice (de Moura e Dias et al., 2023). In contrast, Venturini and colleagues observed increased levels of *Lactobacillaceae* and *Enterobacteriaceae* as well as decreases in Bacteroidales (Venturini et al., 2021). We observed several conserved responses across both ^HMA^mice and MBRA communities. These included loss of *Ruminococcaceae* (UCG-04 and *Anaerotruncus*)*, Lachnospiraceae* (*Sellimonas*), *Erysipelotrichaceae* (*Holdemania*) and increases in an unclassified *Erysipelotrichaceae* and *Anaerostipes* (*Lachnospiraceae*).

Ciprofloxacin is a fluoroquinolone antibiotic with broad-spectrum activity against Gram-negative and Gram-positive bacteria, that has been widely used in clinical practice (Bush et al., 2020). Dethlefsen and Relman reported individualized responses to ciprofloxacin treatment and long-lasting microbiota disruption following treatment (Dethlefsen and Relman, 2011). Rodriguez-Ruiz and colleagues found that ciprofloxacin treatment led to increased levels of *Bacteroides,* several genera of *Lachnospiraceae* (*Blautia*, *Roseburia, Faecalicatena, Dorea*), *Clostridium* (*Clostridiaceae*), and *Eubacterium* (*Eubacteriaceae*), but that levels of *Alistipes* (Bacteroidota), *Faecalibacterium* (Bacillota), and *Coprococcus* (Bacillota) declined in a study of 24 patients (Rodriguez-Ruiz et al., 2024). They also observed increases in *Escherichia* and *Akkermansia* in patients undergoing longer term treatment. *Faecalibacterium* species do not efficiently colonize MBRA communities, so loss of this taxa could not be observed. Consistent with their observations, we observed significant declines in *Alistipes* and increases in *Escherichia-Shigella*. We also observed that levels of *Coprococcus* were lower and levels of *Blautia* and *Dorea* were higher in ciprofloxacin-treated communities compared to untreated communities, but these results were not statistically significant. In contrast to their results, we observed significantly lower levels of *Akkermansia* in ciprofloxacin-treated communities compared to controls.

Clindamycin has long been recognized for its ability to disrupt the microbiome, specifically for its role in increasing susceptibility to *Clostridioides difficile* infection (Brown et al., 2013; Zhang et al., 2022). Clindamycin’s pronounced effects on microbiota are likely due to its broad spectrum activity against anaerobic Gram-positive and Gram-negative bacteria (Guay, 2007). Liu and colleagues studied the effects of clindamycin treatment on the microbiota in Swiss Webster ^HMA^mice colonized with one of three different human fecal samples (Liu et al., 2022) and observed that treatment led to loss of ∼33% of rare taxa and increases in the abundance of *Enterobacteriaceae*, but the microbiota recovered rapidly. In contrast, Buffie et al. observed large, long-lasting decreases in diversity from a single dose of clindamycin in the ileum and cecum of C57Bl/6J mice (Buffie et al., 2012). Many taxa were lost, including members of the *Lachnospiraceae*, *Ruminococcaceae*, and *Turicibacter* (all anaerobic Bacillota), as well as the *Barnesiella* (Bacteroidota). In addition, they observed increases in *Enterobacteriaceae*, *Mollicutes, Akkermansia*, and *Blautia*. In contrast, Markey and colleagues reported increased *Bacteroides* and decreased *Akkermansia* in C57Bl/6 mice (Markey et al., 2021). Across the two models described here, ^HMA^mice exhibited larger changes in richness and taxa abundance, although both models demonstrated loss of *Ruminococcaceae, Lachnospiraceae*, and *Bacteroidaceae* genera, as well as increases in *Enterobacteriaceae*.

Doxycycline, a member of the tetracycline family of antibiotics with higher levels of oral absorption and serum half-life, is another broad range antibiotic (Holmes and Charles, 2009). Sun and colleagues examined effects of doxycycline in CaMK2α-tTA mice (Sun et al., 2024) and observed increased *Lactobacillus, Bacteroides, Parabacteroides*, and Pseudomonadota and decreased levels of *Akkermansia* and *Lachnospiraceae*. In C57Bl/6NCrl mice, Boynton and colleagues also observed increased Bacteroidota and Pseudomonadota and decreased Clostridiales (Boynton et al., 2017). In the MBRA model, we observed very few statistically significant differences compared to controls, with lower levels of *Akkermansia*, four *Ruminococcaceae* genera (UCG-003, NK4A214, *Phocea*, and *Anaerotruncus*), and an unclassified Bacteroidales genus.

Fidaxomicin is a narrow spectrum antibiotic used for treatment of *C. difficile* infection (Goldstein et al., 2012). In 23 patients with mild to moderate *C. difficile* infection, Tannock and colleagues observed that treatment with fidaxomicin led to increases in *Clostridial* clusters XIVa (*Lachnospiraceae*) and IV (*Ruminococcaceae*) and *Bifidobacterium*. Nerandzic and colleagues observed increased levels of *Bacteroides* in *C. difficile* patients treated with fidaxomicin (Nerandzic et al., 2012). Similarly, Yamaguchi and colleagues observed increased *Bacteroidota* in C57Bl/6J mice treated with fidaxomicin (Yamaguchi et al., 2020). Across the two models, we observed five *Ruminococcaceae* and one *Lachnospiraceae* genera were lower in fidaxomicin-treated models compared to controls and *Dielma* (*Erysipelotrichaceae*) was higher compared to controls. Consistent with results described above, higher levels of *Bacteroides* was observed in ^HMA^mice treated with fidaxomicin compared to controls.

Imipenem is a broad spectrum antibiotic used to treat severe infections (Rodloff et al., 2006). Characterization of the microbiota by metagenomic sequencing demonstrated few differences in the composition of taxa (Grall et al., 2017). Characterization of the microbiota by culture-based approaches in patients receiving prophylactic imipenem treatment prior to colorectal surgery demonstrated lower levels of culturable bacteria (Kager et al., 1989), as well as lower levels of *Bacteroides*, fusobacteria, bifidobacteria, clostridia, and lactobacilli. Similar declines in *Bacteroides fragilis* group were observed by culture-based approaches in ten patients, which was accompanied by lower levels of enterococci and *Enterobacteriaceae* (Nord et al., 1985). However, another study demonstrated increased levels of *Enterococcus* species in patients being treated with imipenem (Welkon et al., 1986). Similarly to some of the observations described above, we observed lower levels of Bacteroidales and *Ruminococcaceae* genera, although we also observed higher levels of *Collinsella* (Actinomycetota), *Escherichia-Shigella*, and *Hungatella* (*Lachnospiraceae*).

## Notes

https://zenodo.org/records/14728946

